# DNA-SaM, a robust system for large-scale data storage

**DOI:** 10.1101/2024.11.04.621825

**Authors:** Xiaoluo Huang, Yu Wang, Jiaxin Xu, Ziang Nie, Jiaquan Huang, Yaxin Wu, Zhiwei Qin, Junbiao Dai, Yang Wang

## Abstract

DNA data storage offers a viable strategy to address the impending data explosion. Early attempts to harness DNA as a storage medium have encountered scalability limitations, largely due to the complexity of codec algorithms, the generation of biochemically harmful sequences and lack of a robust architecture. We present “DNA-SaM”, a novel system designed for DNA data storage, which achieves linear computational complexity and strict bio-constraint adherence, ensuring high coding efficiency and fidelity. It encoded data at speeds surpassing classic systems by over 2 orders of magnitude, with this superiority changes across various encoding algorithms. Importantly, DNA-SaM effectively eliminates any sequence that could be deleterious to *in vitro* and *in vivo* biochemical processes, including homopolymer runs, tandem repeat motifs, and potential promoter sequences, *etc*. It also involves an advanced DNA data storage architecture that incorporates a two-tiered indexing system and a novel “storage unit” distribution paradigm for large-scale data storage. It is further validated by practical data storage both *in vitro* and *in vivo* with a 100% success rate. Our system is capable of storing data over 10^39^ PB, which marks a critical advancement in the scalability of DNA-based data storage.

## Introduction

The rapid development of human society has fueled an explosive growth in global data. It is estimated that global data volumes will swell to 300 zettabytes (ZB) by 2028 ^1^. This surge underscores the urgent need for novel storage media with high capacities. Recently, DNA has emerged as a highly promising medium for addressing the impending global data storage crisis. With an information density six to seven orders of magnitude greater than traditional storage media, DNA offers unparalleled potential for data storage. Theoretically, one gram of DNA can store up to 215 petabytes of data, making it an excellent candidate for future large-scale data storage^2^. Moreover, DNA exhibits long longevity, exceptional privacy features, and can be readily replicated, further solidifying its potential as a leading medium for future data storage^3-5^.

Despite DNA’s potential as a viable candidate for addressing the exigent demands of large-scale archival storage, current frameworks for DNA-based data storage have not fully encapsulated the critical components required for extensive application. Over the past decade, pioneering research has brought the concept of “DNA data storage” closer to reality. The early system proposed by Church *et al*. ^6^ employed a simple mapping to translate binary information into A/T/C/G sequences, with 0 corresponding to A or C, and 1 to T or G ^6^. Subsequently, Goldman *et al*. utilized a rotating code for data conversion and introduced a four-fold redundancy to prevent data loss ^7^. However, these early endeavors were limited by low coding efficiency and inadequate management of sequence features that could impair *in vitro* biochemical processes. Notably, the GC content and the prevalence of homopolymers in DNA sequences are known to influence synthesis, sequencing, and amplification efficiencies. The “DNA Fountain” algorithm represents a significant leap forward by optimizing GC content and homopolymer lengths through a sophisticated coding scheme inspired by “Fountain” coding ^2^. This innovation achieved a coding density of nearly 2 bits per nucleotide, marking a key milestone for the feasibility of DNA data storage. Similarly, systems like “Yin-Yang” by Ping *et al*. and “Wukong” by Huang *et al*. encode DNA sequences with high density while maintaining acceptable GC content and homopolymer lengths ^8,9^. To enhance the accuracy of DNA data storage, these works also implement error correction strategies, such as Reed-Solomon (RS) codes ^10^. A recent work, named “DNA-Aeon”, integrates the benefits of “RS code”, “arithmetic code” and “Fountain code” to achieve a high level of data error correction ^11^. Despite these advancements, significant challenges remain existed. Firstly, the computational complexity of these methods is substantial—encoding 1 terabyte (TB) of data could potentially take over one year in a standard computational environment (**Table S8**). Secondly, while GC content and homopolymer lengths are better controlled, other sequence motifs, such as tandem repeats (potentially affecting DNA assembly *in vitro*) and regulatory elements (potentially influencing long-term storage *in vivo*), still lack robust control strategy. Finally, the absence of scalable systems designed for Petabyte-level data storage represents a notable gap in the field. Existing systems have only been validated at the gigabyte scale through a combination of computational simulations and biological experimentation. Together, while DNA data storage holds immense potential to mitigate future data storage challenges, its large-scale implementation requires the development of more robust and scalable systems that overcome the current limitations in sequence optimization and data management.

In this study, we present the “DNA-SaM” system, a novel approach featuring a well-designed codec algorithm and an architecture, that enables the storage of data up to 10^39^ Petabytes (PB). This system excels in encoding data with both high density and fidelity, while also addressing any known sequence-based biological constraints, making it more robust for large-scale data storage applications compared to previous systems. Significantly, the DNA-SaM system exhibits a linear time complexity of O(n), which allows for data encoding at a rate that is two to three orders of magnitude faster than existing systems, within a standard computational framework. With its robust architecture for large-scale DNA data storage, the DNA-SaM system emerges as a promising solution to meet the burgeoning demands of global data proliferation.

## Result and Discussion

### The General Principle of “DNA-SaM” System

Early DNA data storage systems, such as “DNA Fountain”, “Yin-Yang”, “Wukong”, and “DNA Aeon”, aim to encode DNA sequences that are compatible with downstream biochemical processes while maintaining high coding density. However, these systems have largely overlooked the critical necessity of controlling algorithmic complexity. As a result, encoding even 1 TB of data could potentially take several years using current Python programs in a typical office computer environment (**Table S8**). Although DNA data storage is touted as a means to store future large-scale data, this approach is inadequate for meeting the demands of future data storage. To address this gap, we devised “DNA-SaM” (**Fig. 1a**), a system with linear complexity that incorporates data codec, error correction, replacement of forbidden sequences, and a file architecture tailored for large-scale DNA data storage.

**Fig. 1.**
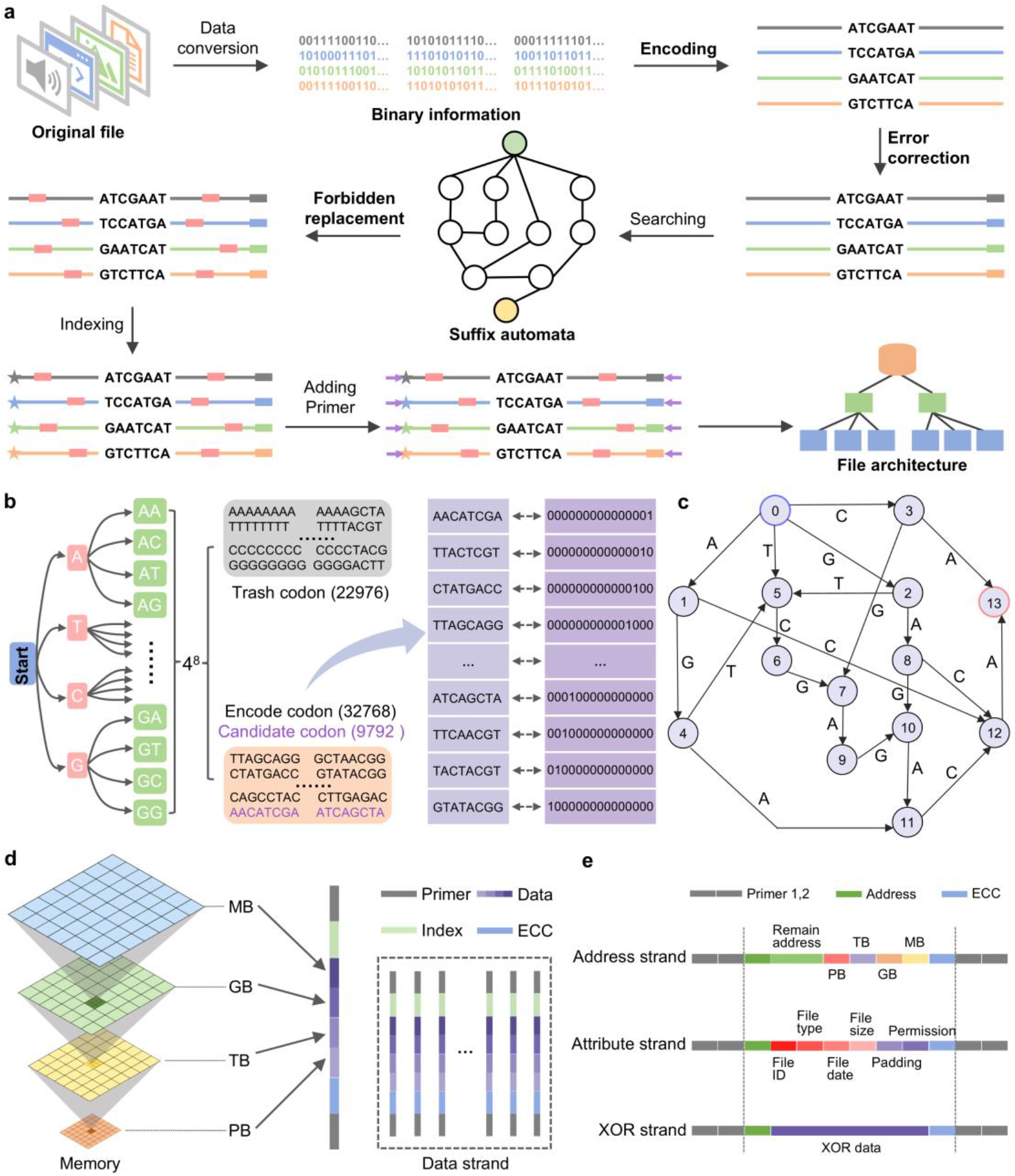
Schematic illustration of the algorithm. **(a)** Overview of the algorithm. **(b)** Codebook generation for mapping-based DNA encoding. **(c)** Illustration of Suffix Automation construction from the sequence “AGTCGAGACA”. **(d)** Segmentation of PB large volume of data and the structure of data strand in for DNA data storage. **(e)** Addition of three extraction strands including address strand, attribute strand, and XOR strand for data storage and retrieval.

#### Generation of a standard Codebook for data codec

Drawing inspiration from the concept of “American Standard Code for Information Interchange (ASCII)” or “Unicode” for text conversion ^12,13^, we first constructed a “Codebook” to facilitate the conversion between digital binary data and DNA sequences. Binary data is encoded into DNA sequences by directly mapping 15 bits to an 8-nucleotide (nt) DNA codon. A detailed description of the mapping strategy is provided in **Supplementary Note 1**, and the process for generating the codebook is illustrated in **Fig. 1b**. To construct the codebook, random combinations of A, T, C, and G are generated *in silico*, resulting in a library of 65,536 8-nt DNA fragments. In line with commercial DNA synthesis standards, which state that GC content above 70% or below 30% can be challenging to synthesize, and considering that most DNA sequencing prefer homopolymer lengths shorter than 4 nt, we established the following constraints: GC content (<62.5%, >37.5%) and homopolymer length (<4 nt). These criteria were used to screen the library, yielding 42,560 valid DNA fragments. The codebook is further divided into two sections: codebook I contains 37,768 DNA codewords for data encoding, and the remaining codewords reserved as potential replacements for forbidden sequences, as discussed below.

#### Error-correction with linearized complexity

While DNA synthesis, PCR amplification, and sequencing are prone to errors, it is essential to implement robust error-correction strategies for accurate DNA data storage ^4,14,15^. Early DNA data storage systems employed the RS code for error correction, which, however, entails significant computational complexity. Given that DNA data storage relies on millions of short synthetic DNA fragments for data encoding, this approach leads to substantial computational resource usage and time consumption. To address this challenge, we have developed an error-correction strategy that integrates the benefits of a previously established single-base error correction method by Cai et al. ^16^ with XOR-based error correction rules. Specifically, Cai’s error correction codes depend on using Varshamov-Tenengolts (VT) codes, and address single-base insertion, deletion, and substitution errors with a linearized complexity. In our standard working flow, four data strings were XORed to produce an additional string, resulting in a 20% additional sequence redundancy for data encoding (**Fig. S1**). Given that the complexity of Cai’s method is O(n) and the XOR redundancy does not change the overall complexity, the overall complexity of our strategy is determined to be O(n). Following error correction comes the core concept of the algorithm, which is the search for and replacement of forbidden sequence using SAM as explained below.

#### Suffix Automation for sequence optimization

Although the codebook controls homopolymer length, it does not entirely eliminate the occurrence of long homopolymers; e.g. the concatenation of encoded codewords can create new single nucleotide repeats. Additionally, similar to early DNA data storage systems, the use of the codebook alone for data encoding may result in the emergence of other tandem repeats and promoter sequences within the encoded DNA sequences. To mitigate this problem, we have developed a rapid encoding and optimization algorithm based on Suffix Automation (SAM) to effectively remove these detrimental sequences (**Fig. 1c**). Sequence replacement serves as cornerstone in the developed scheme for optimizing and managing sequence motifs, which substitutes forbidden sequences, eliminates repetitions, and processes specific motifs using SAM. During the replacement process, SAM is constructed for the target sequence, allowing for the acquisition of each state and the corresponding state transition table, along with the calculation of internal parameters to identify the index of the detected sequence. Additionally, all forbidden sequences are identified within the SAM. To illustrate the core features of this strategy, the operation process of SAM is depicted in **Fig. 1c**. Initially, the SAM is set to its initial state, and all characters in the forbidden sequence are traversed to determine the subsequent state based on the current character and state. Successful traversal indicates the presence of the forbidden sequence in the sequence being inspected. Conversely, the absence of a transfer target for a character in a specific state during traversal confirms that the forbidden sequence does not exist within the examined sequence. Finally, indexed and primers are added to complete the process, with the file structure displayed at the end of **Fig. 1a** and explained in **Fig. 1d**.

#### Architecture for large-scale data storage

In the era of data proliferation, storing PB-level data has become a pressing requirement for institutions and enterprises, *etc*. ^17,18^. The key advancement in this study is the scalability of the system to accommodate over PB-scale DNA data storage (**Supplementary note 2, Fig. S2**). Drawing from the principles of distributed storage architecture found in silicon systems, we have developed a system that encodes data within distributed “DNA units”. Each DNA unit contains a substantial amount of information, ensuring that the damage to one unit does not impede the recovery of data from others. Despite the typically short length of synthetic DNA fragments, we have implemented a two-tier indexing system to assemble large datasets. Within each DNA unit, multiple strands of DNA define its role within the collective “DNA unit” structure. Specifically, one strand contains the address information necessary for the DNA unit to contribute to the larger dataset, while another strand records file metadata such as “file type,” “file size,” “file date,” and “file permissions.” These two strands are combined through an “XOR” operation to create a new strand, further enhancing data integrity and retrieval efficiency.

**Fig. 1d** illustrates the procedure for PB-sized data storage, where data of PB size is segmented into TB, which are further divided into gigabytes (GB), ultimately resulting in MB files that constitute the PB file (128 nt). After appending index (8 nt) and error correction (17 nt) fragments to each end of data, the final data strands (193 nt) are obtained by adding primers (20 nt) generated with WuKong algorithm ^9^ at both ends. To facilitate effective data retrieval, three additional strands are incorporated into the data oligo pool, including an address strand, a file attribute strand, and their XOR strand. The structure of these strands is depicted in **Fig. 1e**. In specific, the address strand serves as a secondary index of data strand, recording file addresses within the sequence cluster. The attribute strand is designed for documenting the attributes of the stored information, such as file ID, type, and the like, with the designed mapping table for various file types presented in **Table S1**. The XOR strand is generated by performing an XOR operation on the main bodies of the address and attribute strands **(Fig. S3**), which plays a critical role in error correction during data retrieval, as it aids in recovering either the address strand or the attribute strand if one is lost. Consequently, the final oligo pool for storing data consists of numerous data strands, along with the address strand, file attribute strand, and XOR strand. A more detailed explanation on PB-level sequence design is provided in **Supplementary note 3**.

### Robustness Analysis of codec performance by “DNA-SaM” system

#### Sequence optimization by “DNA-SaM”

DNA data storage is appealing increasing attention due to its remarkable attributes, including long retention time, high density, and efficient data retrieval mechanism ^19^. By encoding digital information into the sequences of nucleotides—adenine, cytosine, guanine, and thymine— DNA can store vast amounts of data in a compact form. Its stability allows for the preservation of information over thousands of years, far surpassing traditional storage media. However, data integrity in DNA data storage might be compromised by the presence of specific sequence motifs. The formation of repetitive sequences can lead to challenges during both writing and reading processes, as they may cause errors in the synthesis and sequencing of DNA ^20^. Besides, tandem repeats can introduce ambiguity, making it difficult to accurately interpret the originally stored data (**Fig. S4**). Analogous to that the random insertion of transposon sequences in organisms’ genomes can lead to various types of genetic mutations ^21^, a high frequency of repetitions in DNA-encoded sequences poses challenges for DNA synthesis and sequencing, adversely affecting DNA storage. To demonstrate the validity of the replacement effect brought by SAM through wet experiment, PCR assembly was conducted before and after substitution of the repetitive sequences. As shown in **Fig. S5**, a standard PCR using 8 strands of 1 μM oligonucleotides (**Table S2**) successfully assembled the target sequences after the replacement of forbidden sequences. Besides, a heavy band appears when loading the product resulting from 100 μM source oligonucleotides, indicating partial extension without amplification under overloading condition, while no extension was detected at low loading amount. However, PCR assembly failed to produce target sequence when utilizing original sequences with tandem repeats (**Table S3**), regardless of the various concentrations of input oligonucleotides applied. **Fig. S6** shows the success and failure in PCR assembly with various concentrations ranging from 10^2^ to 10^−4^ μM. The overall test on PCR assembly highlights the importance and effectiveness of the substitution method employed in the proposed methodology.

In addition, a number of studies have illuminated that the incorporation of regulatory motifs within exogenous DNA can cause genetic instability ^22,23^. The presence of promoters within these DNAs may trigger unexpected transcriptional events. Moreover, the Shine-Dalgarno (SD) sequence has the potential to induce unintended translation processes. We propose that in the context of DNA data storage, the presence of such regulatory motifs within the encoded DNA could pose a significant risk. There is a genuine concern that such motifs might lead to the degradation of the stored DNA or the demise of the host organism, thereby adversely affecting the recovery of stored information. To mitigate these risks, careful design of the DNA sequences is essential, emphasizing the requirement for robust encoding strategies that minimize the presence of repetitive elements while maximizing data stability and fidelity. SAM offers functionality for editing genomic sequences, as presented in **Table S4**, where a wide variety of important regulatory sequences in the biological genome can be inserted or replaced using the developed replacement algorithm. A comparison of DNA-SaM with previous algorithms, in terms of their encoding quality across varying file types, is displayed in **Table S5**, demonstrating that DNA-SaM outperforms previous methods. In specific, DNA-SaM can successfully avoid TATA box, SD sequence, GAAT box, and tandem repeats, while DNA-Aeon, Goldman’s algorithm, and YYC exhibit limitations in mitigating one or more of these elements. The superior performance of DNA-SaM can be attributed to the use of SAM, which facilitates forbidden sequence detection and replacement. Overall, we advocate the importance of sequence substitution and the effectiveness of SAM.

Notably, the processing time for replacing during data encoding depends on the number of forbidden sequences and total encode sequences. As shown in **Fig. 2a**, the operation time increases as the forbidden sequences rising from 100 to 900. Besides, the more encode sequences scanned for forbidden sequence substitution, the longer it takes for completing the sequence optimization. In specific, the processing time grows from 8.6 s to 12 s when replacing 100 to 900 forbidden sequences within a dataset of 10,000 sequences. Similarly, replacing 100 forbidden sequences takes only 0.85 s when the encode sequence dataset includes 1,000 strands of DNA sequences. These results indicate that the SAM is efficient for rapid sequence optimization and management. Additionally, as presented in **Fig. 2b and Table S6**, replacing 20 forbidden sequences only takes approximately 5.2 s, with time consumption remaining relatively stable across different lengths of the forbidden sequences. To sum up, these *in silico* simulations demonstrate the robustness of SAM, highlighting its favorable function in various scenarios that requires sequence replacement.

**Fig. 2.**
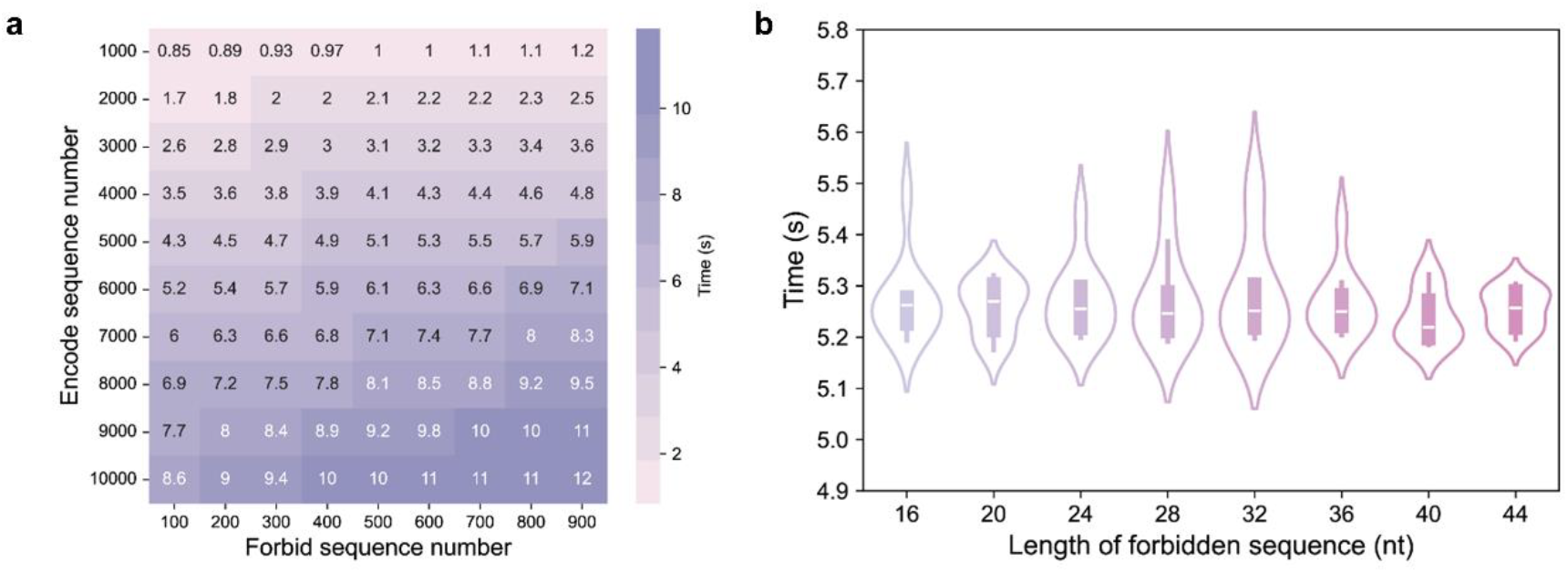
Core part of the proposed algorithm. **(a)** Time consumption for replacing varying numbers of forbidden sequences in different encoded sequence sets. **(b)** Time consumed for dealing with forbidden sequences of varying lengths.

#### Codec Performance of “DNA-SaM”

General biological constraints, such as GC content and homopolymer length, play crucial roles in DNA data storage due to their influence on DNA synthesis, replication, and sequencing. High GC content increases the stability of DNA double helix through stronger hydrogen bonding between G and C, resulting in a higher melting temperature. As a result, strand separation during denaturation, necessary for processes such as PCR amplification or replication, requires greater energy input, either in the form of heat or enzymatic activity. The enhanced stability of GC-rich regions can also hinder the progression DNA polymerase, potentially causing stalling or errors in both natural and synthetic DNA synthesis. Additionally, high GC content increases the likelihood of forming secondary structures, such as hairpins or G-quadruplexes, which can impede DNA synthesis or replication and elevate error rates. Homopolymers present further challenges, as they can cause slippage during synthesis or amplification, leading to unreliable results in DNA data storage. Therefore, optimizing GC content and homopolymer length is critical for the development of robust DNA data storage systems. The developed algorithm is carefully characterized to validate the favorable features equipped for digital information encoding, which is realized through *in silico* simulations by encoding a 10,683-byte text file, a 226,208-byte figure, a 625,899-byte audio file, and a 243,517-byte video. Note that a video of 66,518 bytes was utilized for simulation conducted by YYC, owing to its limitations in encoding a larger video file. As is depicted in **Fig. 3**, the general characteristics are quantified in comparison with major works in terms of homopolymer lengths, GC contents, time consumption, as well as the robustness against mutations. The proposed algorithm generates comparable homopolymers, with lengths within 4 nucleosides, similar to these pioneering works including DNA-Aeon ^11^, DNA Fountain ^2^, the algorithm reported by Goldman *et al*. ^7^, and Yin-Yang codec (YYC) system ^8^ (**Fig. 3a**). Besides, all these algorithms yield GC contents within the range of 30-70% (**Fig. 3b**), ensuring favorable DNA stability for synthesis, storage, and amplification.

**Fig. 3.**
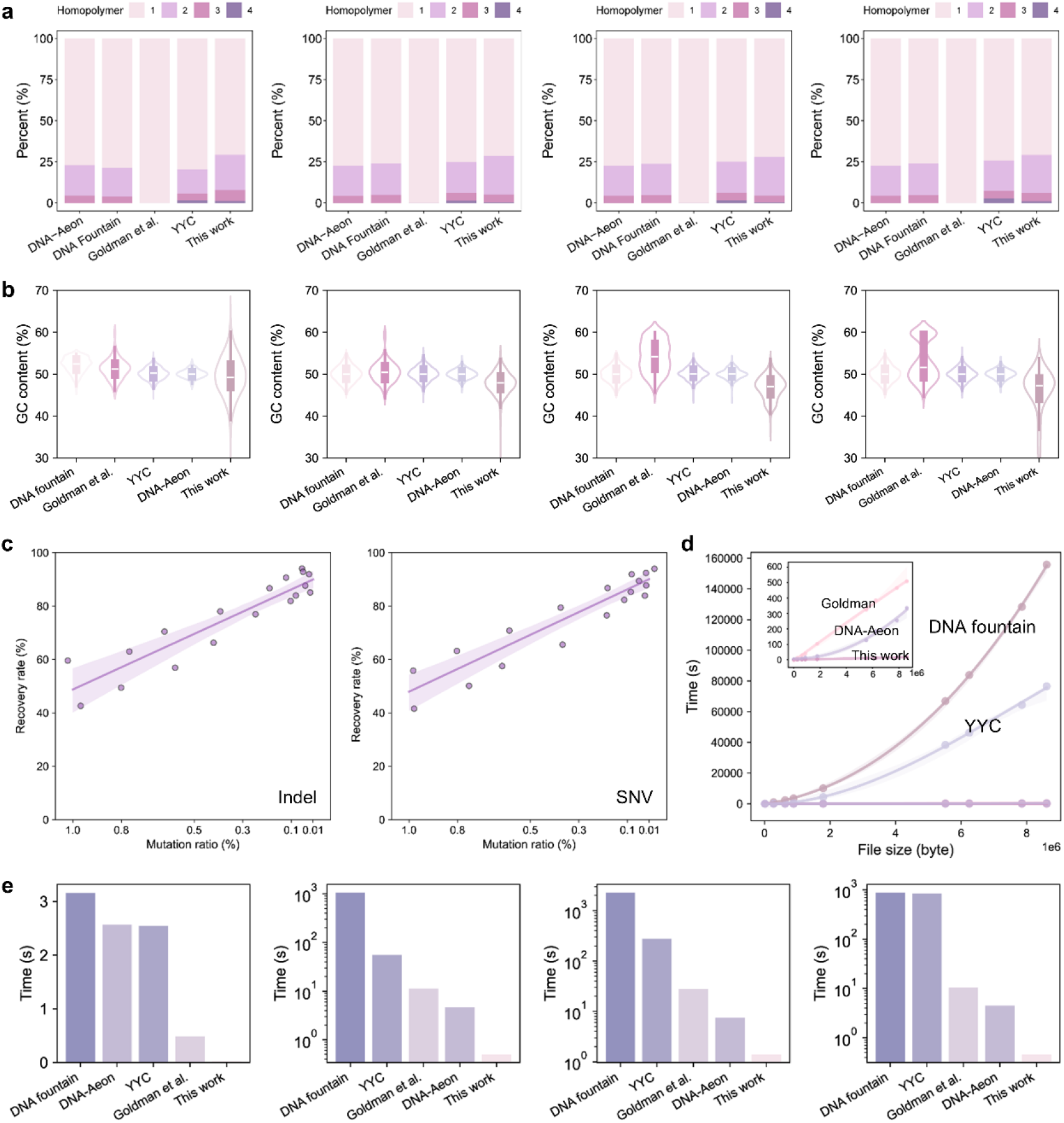
General characteristics of the proposed algorithm for DNA data storage. Comparison of **(a)** homopolymer **(b)** GC uniformity of sequences for text, figure, audio and video files by different methods. **(c)** Robustness characterization of the proposed strategy compared to Yin-Yang code and DNA fountain. **(d)** Time consumed for encoding files differed in size. **(e)** Comparison of time consumption by different DNA data storage schemes for encoding various files. The computer environment: CPU:Inter® Core ™ i7-10700 CPU @2.90 GHZ, RAM, 16 GB, at a Microsoft X64 system.

When exploiting DNA as a storage medium, additional factors such as the robustness against mutations and time efficiency for encoding require careful consideration, as they are decisive to the system’s overall efficiency, reliability, and practicality. To enhance the robustness of DNA data storage against mutations, researchers employ error-correcting codes that introduces redundancy to the data, allowing for the detection and correction of errors ^9,24,25^. Herein, the robustness against mutations ensured by the proposed strategy is confirmed, including insertions and deletions (indels) and single nucleotide variations (SNVs). As shown in **Fig. 3c**, the newly proposed approach in this work exhibits linear data retrieval performance. Our method outperforms the popular DNA fountain code (**Fig. S7**), with the data recovery rate approaching approximately 100% from about 50% when the mutation ratio decreases from 1% to 0.01%. While a trade-off is generally anticipated between robustness and storage density, our method does not sacrifice the storage density while achieving fast encoding speed.

The complexity of the encoding algorithm significantly impacts the time required for efficient data storage, especially in managing error correction, avoiding homopolymers, and maintaining sequence diversity ^26^. To demonstrate the linearized complexity of DNA-SaM, time consumption required to encode files of various sizes is evaluated across different encoding strategies to compare the encoding performance between DNA-SaM and the previous works.

As depicted in **Fig. 3d**, the time consumption for DNA fountain and YYC increases exponentially as the file size becomes larger, indicating nonlinear complexity of these algorithms. DNA-Aeon, and the algorithm reported by Goldman et al., as well as our method, demonstrate faster encoding speeds compared to DNA fountain and YYC, as highlighted by the inset in **Fig. 3d**. Although DNA-Aeon exhibits exponential complexity, it still outperforms Goldman’s method when encoding the file with size within 10^6^ bytes. Similar to Goldman *et al*.’s approach, DNA-SaM encodes data with a time consumption increasing linearly as the size of data increases. Combining the logic of DNA-SaM, this should enable it to have a linear complexity of O(n). In addition, our method achieves 2 to 12 orders of magnitude faster encoding speeds than other approaches presented in **Fig. 3d** when encoding files sized in GB, TB, and PB level (**Supplementary Note 4, Table S7, Table S8**). In specific, DNA-SaM offers at least 2 orders of magnitude faster encoding speed than other algorithms for all the simulated cases, with an exciting superiority for over 12 orders of magnitude than DNA fountain when encoding files sized in PB level. Finally, time consumption for encoding the selected text, image, audio, and video files is exhibited in **Fig. 3e**, demonstrating that the algorithm proposed in this work offers significantly faster encoding speed compared to previous works. It is worth noting that the faster encoding speed for text is invisible in the figure here due to the exponential increase in operational speed relative to other approaches. The novel DNA-SaM algorithm enables completing encoding of the aforementioned files within 1.41 s, with the 10,683-byte text file encoded in just 0.022 s. Overall, the newly proposed work demonstrates favorable performance for information encoding involved in DNA data storage, with demonstrating linearized complexity across various types and sizes of files.

### *In vitro* experimental validation of the proposed algorithm

#### Robustness of data recovery with low molecule copy number

To assess the compatibility of DNA-SaM with practical data storage, three “DNA units”, named “Chest”, “PDF”, and “Star”, each containing around ∼10^4^ DNA strands, were synthesized and selected for proof-of-concept experiments (**Table S9, Table S10**). These oligo pools were objected to standard PCR procedure and sequenced via NGS. As presented in **Fig. 4a**, the oligo pool, containing approximately 10^4^ data strands, an address strand, an attribute strand, and an XOR strand, was dissolved in nuclease-free water to receive a mater pool with a concentration of 10^6^ molecules per microliter. The master pool was further diluted to obtain oligo pools with average copy numbers per oligo ranging from 10^6^ to 10^0^. Serial dilution PCR was performed for each oligo pool, with size verification conducted by gel electrophoresis demonstrating the potential of amplification of each pool (**Fig. S8**). NGS was then performed to assess the robustness of data recovery at low molecule copy numbers. As indicated in **Fig. 4b**, full data recovery was achieved from the amplified fragments originating from sources with molecular concentration of 10^5^ and 10^6^ copies per oligo. However, varying degrees of data loss were observed at lower molecule concentrations, although error correction improved the data recovery rate to some extent. For all oligo pools tested, the raw recovery rate exceeded 98.69% for samples with concentrations higher than 10^5^ copies per oligo, achieving 100% recovery after applying error correction embedded in DNA-SaM. Error correcting codes significantly enhanced the data recovery rates, such as improving the ratio from 20.18%, 34.17%, and 76.29% to 36.51%, 51.95%, and 91.81% for the oligo pools encoding distinct information, that is, “Chest”, “PDF”, and “Star”, at molecular concentration of 10^4^, respectively. In conclusion, DNA-SaM exhibits robust performance for data storage, with reliable data recovery across a range of molecular concentrations.

**Fig. 4.**
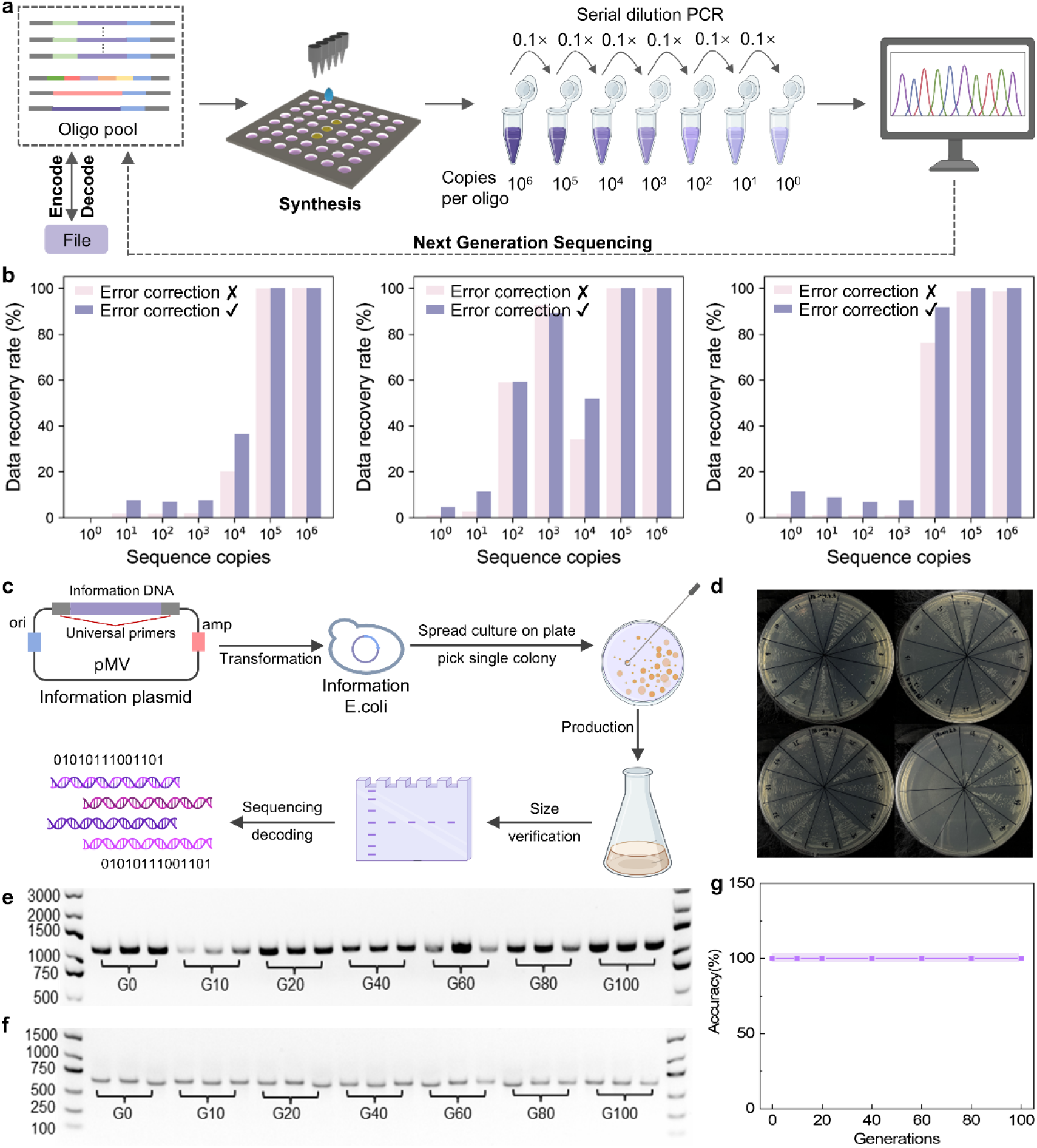
Experimental validation of the proposed method. **(a)** *In vitro* evaluation by analyzing data recovery rates after series dilution PCR. Structure of file-encoded sequences were presented in the box. **(b)** Data recovery rate for three files (“Chest”, “PDF”, and “Star”) under various molecular concentrations ranging from 10^6^ to 10^0^ copies per oligo. **(c)** Workflow of *in vivo* data storage validation. **(d)** Colony formation of information-inserted *E*.*coli*. **(e)** Identification of information plasmids and **(f)** information sequences after serial passaging ranging from 0 to 100 generations. **(g)** Data recovery from *E*.*coli*. after serial passaging until 100 generations. BR: before replacement. AR: after replacement. BR1 and BR 2 represent for performing PCR assembly with oligonucleotides of 1 μM and 100 μM concentrations. G0 to G100: 0 generation to 100 generation.

### *In vivo* experimental validation of the proposed algorithm

*In vivo* DNA data storage demonstrates significant advantages due to its potential for rapid, stable, and cost-effective data replication and retrieval. Gene sequence is suitable for *in vivo* storage and are commonly employed to store small files that can be transform into bacterial for preservation. Gene are more likely to exhibit special structures, which can affect cell growth and development, and even lead to cell death. The encoding algorithm based on suffix automata in gene storage can avoid harmful sequences such as repetitive sequences, promoter sequences, and special motifs in advance through this algorithm, as shown in **Table S4**, so that the encoded DNA sequence can stably exist and be inherited in the cell.

Using the proposed method, this study encoded “A very short story” by Ernest Miller Hemingway into a total of 41 strands DNA sequences (**Table S11**). Each sequence stores 105 bytes data and has a length about 500 nt. Plasmids were constructed with these DNA sequences to enhance stability, sequencing compatibility, transformation and passaging. The overview of the *in vivo* validation is illustrated in **Fig. 4c**, where information sequence-included plasmids were transformed *E*.*coli* strains, followed by colonies formation on solid culture plate and serial passaging in liquid culture medium after isolating a single colony. Plasmids were harvested from the resulting culture medium at various growth stages, followed by size verification and Sanger sequencing to confirm the successful data storage and access.

Gel electrophoresis was conducted to separate and identify the sizes of plasmids and their PCR products. (**Fig. S9**) offers the size information confirmed by gel electrophoresis for the overall information-encoded plasmids and information sequences, indicating the potential for extracting and replicating plasmids and information-encoded sequences from *E*.*coli*. cells. The stability, bulk replication, and recovery of information were assessed through a standard cell passaging process, followed by size verification and sequencing. Three strains containing different information sequences were randomly selected for the serial passaging experiment. The growth curves, monitored at 15-min intervals, indicated approximately 40 min for one generation passage, as shown in (**Fig. S10**). The cells were cultured continuously to 100 generations with fresh media changed frequently (OD600 of 0.05 to 1.6), with using Glycerol stock solutions as 0 generation. As shown in **Fig. 4d**, cells exhibited robust growth after passaging, suggesting the bulk replication capacity of the bacteria strains. Plasmids were harvested from 0, 10, 20, 40, 60, 80, and 100 generations and analyzed by gel electrophoresis (**Fig. 4e, f**) before and after PCR amplification, indicating long-term storage stability and reliable data replication. Furthermore, plasmids from these generations underwent to Sanger sequencing, with readable peaks for DNA sequences collected at each generation presented in (**Fig. S11**) and full alignment of to their designed sequences (**Fig. S12**). Consequently, information retrieval was achieved with 100% accuracy after generation passaging (**Fig. 4g**), validating the reliability and effectiveness of the proposed algorithm for DNA data storage.

### Exploiting the potential PB-level DNA data storage

Algorithm’s scalability is crucial for accessing the capacity of handling increasing data volumes, which determines the feasibility and practicality of large-scale data storage. The developed strategy, involving a collection of DNA units to form an architecture capable of extended data storage, demonstrates unprecedented performance for storage of over 10^39^ PB without compromising its efficiency (**Supplementary note 2, Fig. S2**). This novel architecture for large data storage includes dividing of big data into small units, followed by encoding these units separately. To provide an overview of functionality of the proposed mechanism for PB-level DNA data storage, the theoretical maximum storage capacities for our system are calculated.

As displayed in **Fig. 5a**, the maximum storage capacities increase as the sequence length increases, showing over 5×10^57^ PB capacity when encoding data into 300-nt DNA sequences. As indicated in **Fig. 5b**, the capacities of a single unit increase with the encode sequence length, achieving a unit storage capacity of 300 kB to 550 kB when the encoded sequences’ lengths increase from 200 to 300 nt. These excellent increasement in both unit and maximum storage capacity indicates the huge potential of DNA-SaM for large data storage. Besides, the splitting time and encoding time versus data size are depicted in **Fig. 5c**, with both the splitting time and the encoding time increase with the size of the files. In specific, the time for data separating and encoding are simulated *in silico*, which confirms the successful operation of files ranging from several MB to tens of GB. Based on the observed linear relationship between operation time and data size, we estimate that the splitting time will reach approximately 10^9^ s and the encoding time will be less than 10^11^ s for a 1 EB file.

**Fig. 5.**
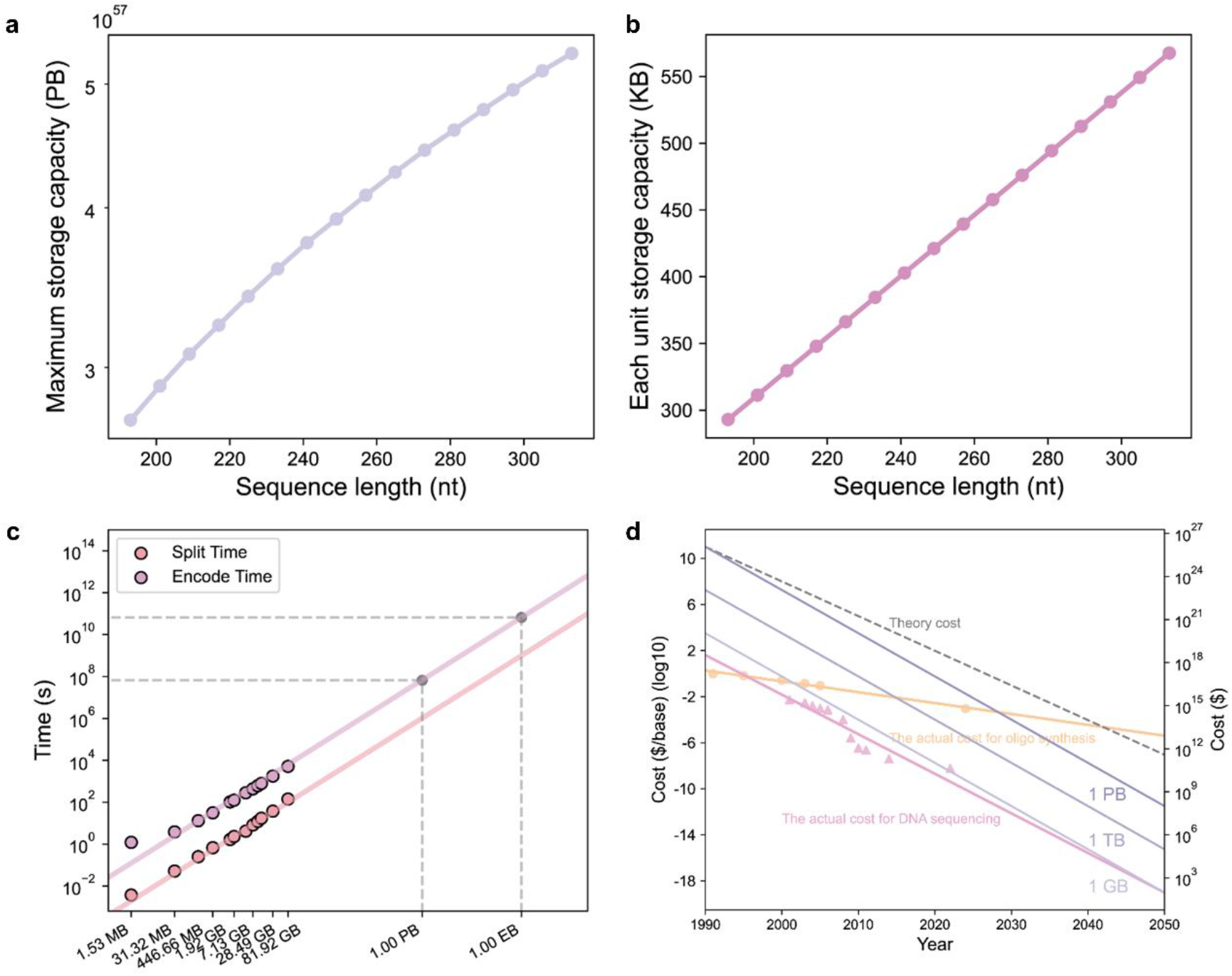
PB data storage in DNA sequences achieved by the proposed algorithm for DNA data storage. **(a)** Increasement of the maximum of the storage capacity based on DNA-SaM following the increase in sequence length. **(b)** Sequence length dependence of unit storage capacity. **(c)** Projection of time consumed for large volume of data encoding. **(d)** Projection of cost for DNA data storage note that the right Y axis is drawn for three purple lines (1 GB, 1 TB, 1 PB), while the left axis is for the other lines.

Cost remains a key factor in extending the practical usage of DNA data storage. Therefore, we further calculated the projected costs for the developed PB-level DNA data storage strategy, using the Moore’s law as a reference. The high expense associated with DNA data storage scheme is mainly resulted from DNA synthesis. As shown in **Fig. 5d**, the synthetic cost per base has gradually decreased from 0.96 $ in 1991 to 9.1×10^−4^ $ in 2024 ^27^. If this trend continues, we project that the cost will drop to 4.00×10^−6^ $ per base by 2050. Unlike the synthetic cost, the cost of DNA sequencing has decreased dramatically due to the advent of NGS, with the price per base falling from 5.3×10^−3^ $ in 2001 to 6×10^−9^ $ in 2022. Using these trends, we simulated the costs for storing 1 GB, 1 TB, and 1PB of data with the proposed encoding strategy.

As indicated by the purple lines in **Fig. 5d**, we predict that the costs for these storage amounts will decrease to 96.01 $, 9.83×10^4^ $, and 1.01×10^8^ $ by 2050, respectively. Given ongoing endeavors by researchers in academia and industry, we believe the cost will continue to decrease, making large-scale DNA data storage a reality in the near future.

## Conclusion

In conclusion, the past one decade has seen the experimental validation of DNA as a viable medium for data storage, with its high density and longevity being successfully demonstrated. Despite the anticipation that DNA data storage will be pivotal for large-scale archiving, current systems grapple with practical challenges that hinder the realization of this goal. We have addressed these challenges through the development of a “DNA-SaM” system that operates with linearized complexity, significantly enhancing the potential for large-scale DNA data storage. Our approach not only accelerates computation times, which can be further optimized with high-performance computing or algorithmic enhancements, but also allows for the customization of sequence constraints. This versatility ensures compatibility with diverse biochemical environments and enables the incorporation of identifying motifs, thereby adding a layer of robustness to the stored data. For example, using DNA-SaM, this study successfully eliminates TATA boxes, SD sequences, GAAT boxes, and tandem repeats—elements that are undesirable in DNA data storage due to their potential to introduce errors during DNA synthesis, sequencing, and data interpretation. With the capability to handle petabyte-scale data storage, our system has the potential to revolutionize the management of large datasets. *In vitro* and *in vivo* validations confirm the efficacy of our architecture. Data is recovered with 100 % accuracy at molecular concentrations above 10^5^ copies per oligo, though significant data loss occurred at concentrations as low as 10^2^ copies per oligo. When inserting the encoded information in plasmids and transforming into *E*.*coli*, data can be replicated easily within a short time period, maintaining 100% data accuracy after passaging across 100 generations. Furthermore, the maximum storage capacity is estimated to be 10^39^ PB by our system. However, the economic barrier of DNA data storage remains a significant obstacle, with costs currently prohibitive for widespread adoption. Looking ahead, we are optimistic that the cost trajectory of DNA synthesis will mirror Moore’s Law, with a projected decrease to 4.00×10^−6^ $ per base by 2050^28,29^. Concurrently, the cost of DNA sequencing has seen a dramatic reduction, a trend that we have leveraged to forecast the future costs of storing data using our encoding strategy. Our simulations suggest that by 2050, the costs for storing 1 GB, 1 TB, and 1 PB of data could be as low as 96.01 $, 9.83×10^4^ $, and 1.01×10^8^ $, respectively. At that time, DNA-SaM should be a cost-effective solution for a broad range of applications, from cloud storage to medical and national archives. Ultimately, the DNA-SaM system stands to significantly advance the field of large-scale data storage in DNA, paving the way for a future where DNA-based storage is not only feasible but also economically viable.

## Materials and Methods

### PCR assembly

Primers was dissolved to 10 μM, with P1 and P2 utilized for PCR assembly before forbidden sequence replacement, while P’1 and P’2 used for that after replacement. All oligonucleotides were dissolved to 10^2^ μM, followed by series dilution to 1 μM, 10^−2^ μM, and 10^−4^ μM. PCR were conducted by preparing a mixture containing 5 μL Q5 reaction buffer, 0.5 μL 10 mM dNTPs, 1.25 μL of 10 μM forward and reverse primers, 1 μL of O1 to O13 (O’1 to O’8 for PCR assembly after forbidden sequence replacement) for each, and 0.25 μL Q5 high-fidelity DNA polymerase, the mixture was brought to 25 μL for PCR reaction. PCR reaction was proceeded with the following procedure: an initial denaturation at 98 °C for 30 s, 30 cycles at 98 °C for 10 s, 56 °C for 30 s, and 72 °C for 30 s, a final extension at 72 °C for 2 min. After the PCR, 1 μL of the PCR product was loaded to the well of 3% agarose gel for electrophoresis at 150 V for 30 min to identify their lengths.

### Dilution PCR for oligo pool

The oligo pools for Star, Chest, and PDF contains 9743, 10651, and 11024 strands of 193 nt oligos. Primers for PCR are summarized in **Table S12**. The received oligo pool was dissolved to 10^6^ molecules per microliter as suggested by the supplier, followed by series dilution to 10^5^, 10^4^, 10^3^, 10^2^, 10^1^, and 10^0^ molecules/μL. PCR were conducted by preparing a mixture containing 10 μL Q5 reaction buffer, 5 μL 2 mM dNTPs, 2.5 μL of 10 μM forward and reverse primers, 1 μL of template solution, 10 μL GC enhancer, and 0.5 μL Q5 high-fidelity DNA polymerase, the mixture was brought to 50 μL for PCR reaction. PCR reaction was proceeded with the following procedure: an initial denaturation at 98 °C for 30 s, 30 cycles at 98 °C for 10 s, 56 °C for 30 s, and 72 °C for 30 s, a final extension at 72 °C for 2 min. The experiment was grouped into two, with no GC enhancer added for group 1. After the PCR, 1 μL of the PCR product was loaded to the well of 3% agarose gel for electrophoresis to identify their lengths.

## Acknowledgements

This study was supported by National Key research and Development Program of China (2021YFF1201700), National Natural Science Foundation of China (32201207), Shenzhen Science and Technology Program (KQTD20180413181837372, RCYX20221008092950122), Innovation Program of Chinese Academy of Agricultural Sciences and Shenzhen Outstanding Talents Training Fund.

## Author contributions

Yang. W proposed the idea of introducing the “*Suffix Automation*” to DNA-SaM system. X. H, Yang. W, Z. N developed the essential codec methods and architecture of DNA-SaM system.

Z. N wrote the initial codes of “DNA-SaM” system. Yu. W optimized the architecutre of DNA-SaM system and wrote the final codes of “DNA-SaM” system. X. H, Yu. W & J. X designed the experiments, figures and tables. Yu. W, J. X, Z. N, J. H& YX. W carried out computational or biological experiments to validate the system, prepared the figures and tables for the paper.

J. X, X. H & Yu.W wrote the initial draft of the paper. X. H, J. X, Yang. W, Z. Q & J. D polished and finalized the paper. X. H, Yang. W, J. D & Z. Q coordinated the resources and supervised the study. All authors read and approved the final draft of the paper.

## Competing interests

Z. N, Yang. W and X. H, et.c applied two patents about DNA-SaM system with the NO. CN202311653624.9, CN202410291695.7.

## Notes

### Competing Interest Statement

Z. N, Yang. W and X. H, etc. applied two patents about DNA-SaM system with the NO. CN202311653624.9, CN202410291695.7.

