## Supplementary material 1 for "DNA-SaM, a robust system for large-scale data storage"

**Table of Contents**

**Supplementary Notes**	4

Supplementary Note 1. Details of Encoding and Decoding Algorithm	4

Codebook Generation	4

Sequence Optimization	5

Decoding Process	6

Supplementary Note 2. Scalability Calculation for Large-data Storage	8

Supplementary Note 3. Details of Storage System	9

File System	9

Sequence Structure	9

Supplementary Note 4. Time prediction when encoding files with different sizes by varied algorithms including Goldman *et al.*, YYC, DNA fountain, DNA-Aeon, and DNA-SaM (This work)	11

**Supplementary Figures**	12

Fig. S1. Illustration of XOR error correction strand generation.	12

Fig. S2. Illustration of file structure for scalability calculation.	12

Fig. S3. The formation of XOR strand.	12

Fig. S4. Examples of tandem repeats formation in the sequences produced by algorithms proposed by Goldman *et al.* ^1^ and Yin-Yang codec system ^2^.	13

Fig. S5. In vitro PCR assembly that demonstrates the importance of substitution.	14

Fig. S6. PCR assembly before and after forbidden sequences replacement under varying concentrations of oligonucleotides ranging from 10^2^ to 10^-4^ μM.	14

Fig. S7. Robustness characterization of DNA-SaM compared to Yin-Yang code and DNA fountain.	14

Fig. S8. Size verification after serial dilution PCR for oligo pools synthesized for three files. The oligo pools were diluted from 10^6^ to 10^0^ molecule copy numbers.	15

Fig. S9. Size verification performed by gel electrophoresis. (a) Verification of 41 strands of information-encoded DNA sequences inserted in plasmids. (b) Verification of 41 strands of information-encoded DNA sequences.	15

Fig. S10. Growth curves of three randomly selected *E.coli* strains containing information-encoded DNA sequences.	15

Fig. S11. Sequencing result for randomly selected three strains after culturing 0 and 100 generations.	16

Fig. S12. Sequence alignment for the randomly selected three strains after culturing 0, 10, 20, 40, 60, 80 and 100 generation.	34

**Supplementary Tables**	37

Table S1. File type encoding table.	37

Table S2. DNA sequences used for PCR assembly after replacement.	38

Table S3. DNA sequences used for PCR assembly before replacement.	39

Table S4. Editing of bio-functional sequences in genome by SAM.	40

Table S5. Comparison of performance guaranteed by various algorithms when encoding text, image, audio, and video files.	41

Table S6. Operation time and times of forbidden sequence replacement for various numbers and types of sequences.	41

Table S7. Time consumption for encoding various sizes of data by algorithms including Goldman *et al.*, YYC, DNA fountain, DNA-Aeon, and this work. Unit of time: second.	42

Table S8. Time consumption for encoding files in GB, TB, and PB level by algorithms including Goldman *et al.*, YYC, DNA fountain, DNA-Aeon, and this work. Unit of time: second.	42

Table S9. File encoding details for experimental verification.	43

Table S10. Oligo pools designed for proof-of-concept experiments.	44

Table S11. DNA sequences designed for storing A very short story by Ernest Miller Hemingway.	44

Table S12. Primers for PCR amplification of oligo pools generated from Star, Chest, and PDF	44

**Supplementary Notes**

**Supplementary Note 1. Details of encoding and decoding algorithm**

**Codebook Generation**

To expedite the encoding process, a direct mapping codebook is employed. The codebook maintains a fixed binary length of 15 bits, while the DNA sequences consist of 8 nucleotides. The designed codebook ensures a GC content of 37.5-62.5% and a homopolymer length of 4 or fewer nucleotides, which helps guarantee the accuracy of synthesis and sequencing processes included in DNA data storage systems. The algorithm to generate a codebook is displayed as the follows.

**Algorithm 1** DNA codon design

**Input:** the length of original DNA codon *l_0_*, max length of homopolymers in DNA codon *l_1_*, the upper limit of GC content 𝑟_𝑚𝑎𝑥_, the lower limit of GC content 𝑟_𝑚𝑖𝑛_.

1: **Initialization**: depth_𝑖𝑛𝑖_ ← 0，currentSeq_𝑖𝑛𝑖_ ← null，continuity_𝑖𝑛𝑖_ ← 0，lastBase_𝑖𝑛𝑖_ ← null，gcContent_𝑖𝑛𝑖_ ← 0，result ← null.

2: **function** GENERATE (depth, currentSeq, continuity, lastBase, gcContent)

3: **if** depth == 𝑙_0_ **then**

4: **if** gcContent / 𝑙0 ≤ 𝑟_𝑚𝑎𝑥_ **and** gcContent / 𝑙0 ≥ 𝑟_𝑚𝑖𝑛_ **and** continuity ≤ ⌊𝑙*_1_* ⌋/2 **then**

5: add currentSeq to result

6: **end if**

7: **return**

8: **end if**

9: **if** depth == ⌊𝑙1 ⌋/2 + 1 **and** continuity == ⌊𝑙1 ⌋/2 + 1 **or** continuity = 𝑙1 **then**

10: **return**

11: **end if**

12: **for** i **in** {A, T, G, C} **do**

13: **if** i == lastBase **then**

14: continuity ← continuity +

15: **else**

16: continuity ← 1

17: **end if**

18: Concatenate i to currentSeq

19: **if** i == G **or** i == C **then**

20: gcContent ← gcContent +

21: **end if**

22: **end for**

23: GENERATE (depth + 1, currentSeq, continuity, i, gcContent)

24: **end function**

25:

26: GENERATE (depth_𝑖𝑛𝑖_, currentSeq_𝑖𝑛𝑖_, continuity_𝑖𝑛𝑖_, null, gcContent_𝑖𝑛𝑖_)

**Output:** result of DNA codon set

**Sequence Optimization**

A rapid optimization encoding algorithm for DNA based on suffix automata consists of three steps: direct information encoding, forbidden sequence replacement, and error correction code addition. Firstly, in direct information encoding, data is converted from binary form to DNA sequences using a pre-designed mapping table. Next, forbidden sequences are substituted utilizing a classic suffix automaton data structure. By constructing a suffix automaton for the DNA sequence, forbidden sequences can be identified. For instance, in the DNA sequence "AGTCGAGACA" with the forbidden sequence "TCGA," a suffix automaton with 14 states is generated. The search starts from state 0, transiting through state 5 with character T, state 6 with character C, state 7 with character G, and ending at state 9 with character A, indicating the forbidden sequence is found at position 2. If no complete path exists for the forbidden sequence in the automaton, it indicates the absence of that sequence in the DNA.

**The rapid DNA optimization encoding algorithm based on the suffix automaton consists of the following steps:**

1. Binary Data Reading: Read the binary data from the file, denoted as *B*, and divide *B* into blocks of 30 bytes, denoted as *b_i_*.

2. Block Encoding: Encode each block *bi* using a direct mapping table to convert bi into *d_i_*, where every 15 bits are encoded into an 8-base DNA sequence block, and each DNA sequence block *d_i_* is 128 bases long.

3. Forbidden Sequence Replacement: Replace forbidden sequences in *di* and record the replaced sequences and their indices, resulting in *r_i_*.

4. Single-base Error Correction Code Addition: Add single-base error correction codes to *ri*. After error correction encoding, each sequence has an additional 17-base redundant error correction code at the end, converting *r_i_* to *e_i_* with a length of 145 bases.

5. Address Block Addition: Add an address block to the head of each sequence, encoding its sequence number as an 8-base sequence according to its division order, converting *e_i_* to *a_i_* with a length of 153 bases.

6. Adding Specific Primers: Add 20-base specific primers p1 and p2 to both ends of each *a_i_* sequence, resulting in a final sequence length of 193 bases.

**Decoding Process**

The decoding process is the inverse of the encoding process:

1. Sequencing Data Assembly: Start with the original sequencing data and assemble it.

2. Specific Primer Matching: Match the assembled sequences with specific primers and retain sequences with complete specific primers.

3. Sequence Segmentation: Split the filtered sequences into address blocks, data blocks, and error correction codes.

4. Error Correction Recovery: Correct the data blocks and error correction blocks. Correctly recovered sequences are considered correct, and the address information of unrecoverable sequences is recorded.

5. Consistency Alignment: Perform consistency alignment on sequences with the same address block that failed error correction to generate a consensus sequence for secondary error correction recovery.

6. XOR Recovery: For sequences that failed secondary error correction, perform XOR recovery by XOR continuous sequences containing the failed sequence to generate XOR supplements.

7. Direct Mapping Decoding: Decode all successfully error-corrected sequences using the direct mapping table, converting DNA sequences to binary sequences.

8. Final Decoded File Generation: Concatenate all sequences according to the address block order to generate the final decoded file.

**Supplementary Note 2. Scalability Calculation for Large-data Storage**

The structure of the secondary index chain is similar to that of data strand, except for that the encoded data is replaced by address units. As shown in **Fig. S2**, a data strand contains specific primers for the encoded files at each end, an 8-nt address storing the total address of the information colony, a 17-nt error correction code, and a 128-nt data segment. Given that the DNA codebook in DNA-SaM comprises clusters of 8-nt codes, we also use 8-nt as an address segment. Therefore, the secondary index strand includes addresses of MB, GB, TB, and so forth, indicating the location of the colony, specifically, in which MB, GB, and TB the colony exists. Since the total encoding data block is 128 nt long and each address unit is 8 nt long, up to 16 address units can be accommodated. Each subsequent unit (after MB, GB, and TB) is 2^10^ times larger than the previous one, meaning that the maximum data capacity for this secondary storage structure is 2^160^ MB (2^130^ PB), or approximately 10^39^ PB of data.

**Supplementary Note 3. Details of Storage System**

**File System**

In a computer management system, information is systematically organized into files rather than being stored arbitrarily. A file-based information storage system is essential for efficient data processing. Leveraging the characteristics of DNA information storage, we designed two DNA file storage systems: one for *in vivo* applications and another for *in vitro* applications.

The file system utilizes a two-level indexing method to connect file addresses with sequence indices within the files. As illustrated in **Fig. 1**, the PB memory is segmented into 1024 TB, each TB into 1024 GB, and so forth, with the smallest unit being MB. The address strand in the DNA sequence cluster holds the corresponding indices for PB, TB, GB, and MB.

**Sequence Structure**

The structures of sequences are displayed in **Fig. 1**, **Fig. S2** and **Fig. S3**. The address strand functions as a secondary index alongside the data strand index, primarily recording file addresses within the sequence cluster. Each end features two pairs of specific primers with length of 20 nt: file-specific primers and address strand-specific primers. Inward from both ends, there is an 8 nt address block that records its position within the sequence cluster, followed by a 17 nt error correction code at the tail end. There is a 56 nt reserved block for future memory expansion, followed by four 8 nt memory blocks representing PB, TB, GB, and MB.

To document file attributes, a file attribute strand was designed. This strand incorporates two pairs of primers, with the file-specific primer at the end matching the file data strand and the adjacent attribute strand-specific primer used to distinguish and extract this strand. The address block and error correction code are consistent with those in the data block.

The file ID block comprises two 8 nt base segments, each recording a digit. Each file ID is represented by these two numbers, uniquely indicating the filename through a relational table, with a total length of 16 nt. The file type block records the file type, currently supporting text, audio, video, compressed files, etc., with different blocks corresponding to different file extensions, 4 nt in length (**Table S1**). The timestamp block records the last modification time of the file, 24 nt in length. The file size block records the memory size occupied by the file, 20 nt in length. A padding block is included to equalize the length with the data strand, measuring 4 nt in length. The file permissions block records file usage permissions, with types such as 'user', 'admin', and 'other', with 20 nt in length. The total length of the file attribute strand is 193 nt.

Finally, the XOR strand is used to recover either the address strand or the attribute strand if one is lost. The primers, address block, and error correction code are the same as the other two strands, with the remaining 88 nt region internally generated by XORing the same positions from the other two strands.

**Supplementary Note 4. Time prediction when encoding files with different sizes by varied algorithms including Goldman *et al.*, YYC, DNA fountain, DNA-Aeon, and DNA-SaM (This work).**

Based on the time consumption for encoding various files by Goldman *et al*.^1^, YYC ^2^, DNA fountain ^3^, DNA-Aeon ^4^, and DNA-SaM by simulations *in silico* (**Table S7, Fig. 3d**), lines representing performances of each algorithm were undergone a linear regression fitting by Linear Regression Method proposed by Selearn. If we define S as file size, T as the required encoding time, the equations for each algorithm are as follows:

For Goldman *et al.*:

T = 6.04 × 10^-5^ S – 6.027

For YYC:

T = 3.26 × 10^-4^ S^2^ + 6.027 × 10^-11^ S – 164.85

For DNA fountain:

T = 2.41 × 10^-3^ S^2^ + 1.78 × 10^-9^ S + 56.11

For DNA-Aeon:

T = -5.06 × 10^-7^ S^2^ + 4.35 × 10^-12^ S + 6.81

This work*.*:

T = 1.89 × 10^-6^ S + 0.128

We projected the time consumption for encoding files with sizes of GB, TB, and PB following these equations above, and summarized data in **Table S8**. It indicates that DNA-SaM displays at least 2 orders of magnitude faster encoding speed than other algorithms, especially demonstrating the superiority for over 12 orders of magnitude than DNA fountain when encoding files sized in PB level.

**Supplementary Figures**

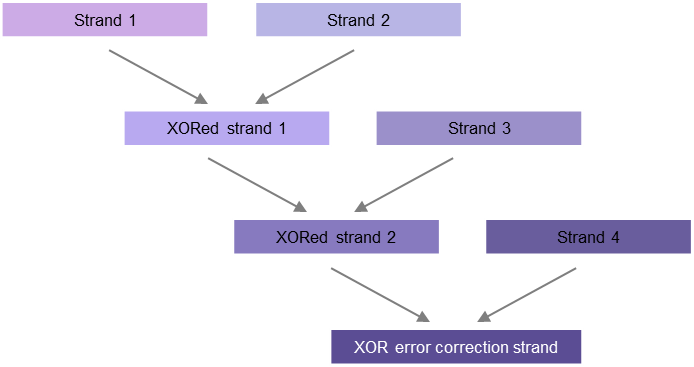

**Fig. S1.** Illustration of XOR error correction strand generation.

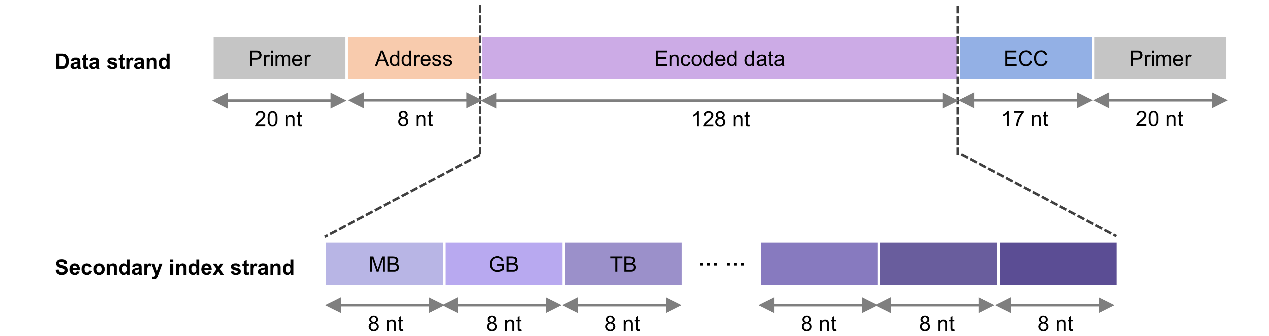

**Fig. S2.** Illustration of file structure for scalability calculation.

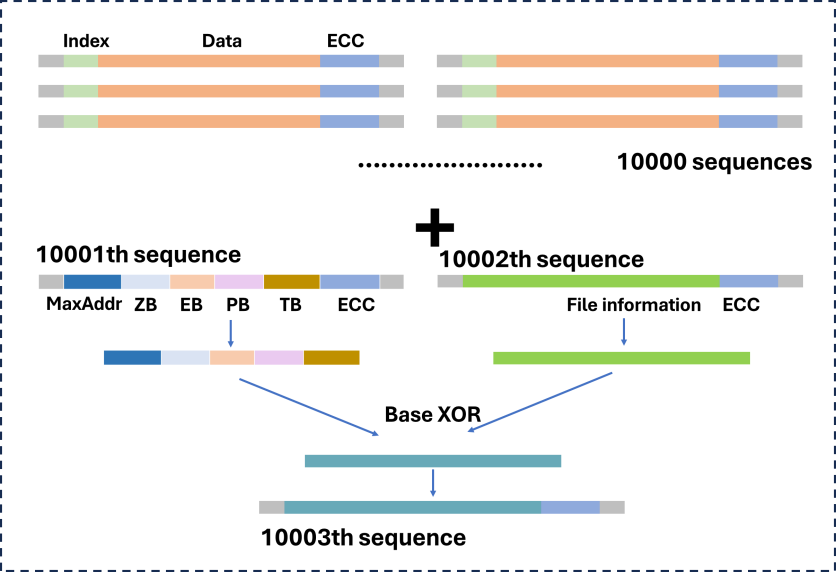

**Fig. S3.** The formation of XOR strand.

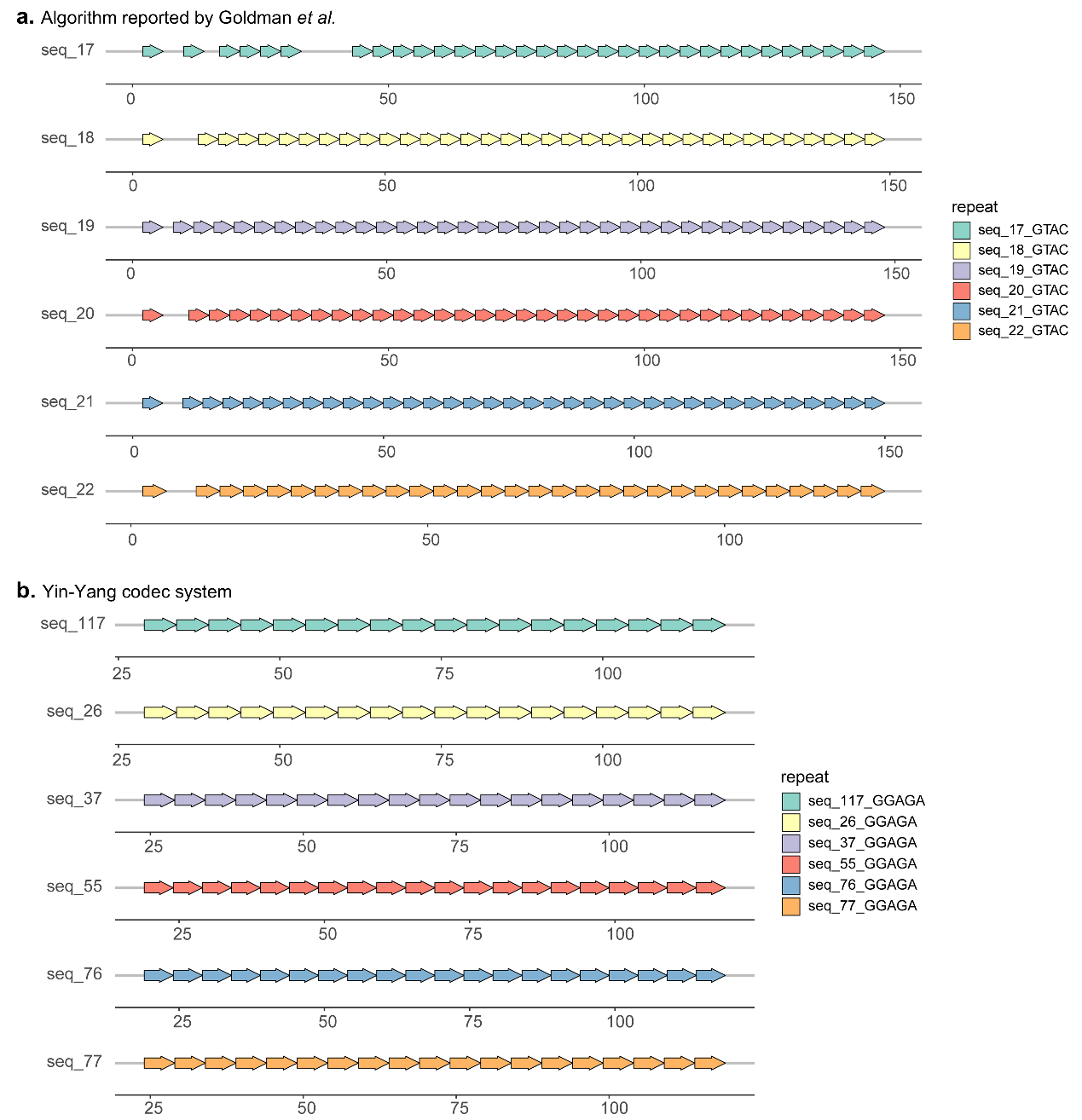

**Fig. S4.** Examples of tandem repeats formation in the sequences produced by algorithms proposed by Goldman *et al.* ^1^ and Yin-Yang codec system ^2^.

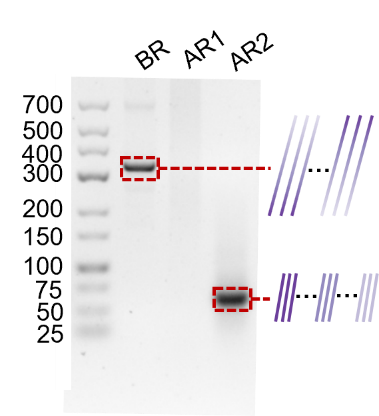

**Fig. S5.** In vitro PCR assembly that demonstrates the importance of substitution.

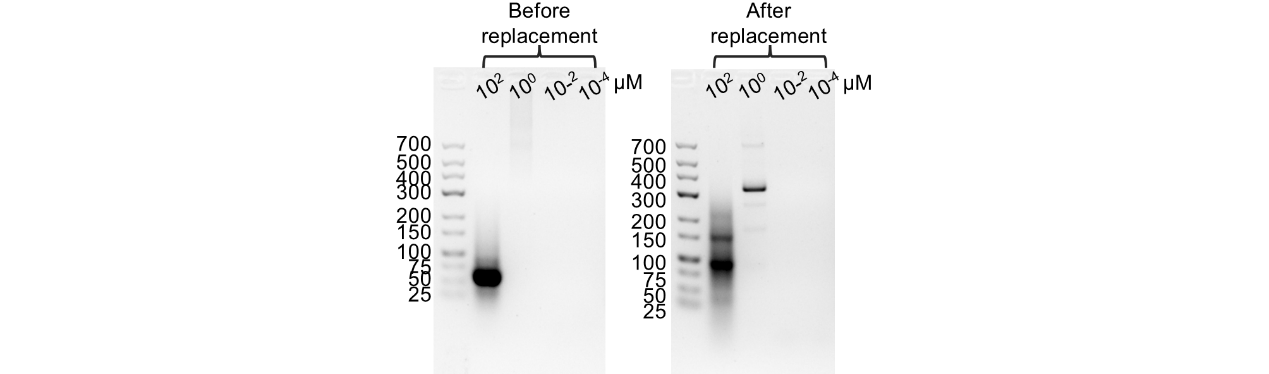

**Fig. S6.** PCR assembly before and after forbidden sequences replacement under varying concentrations of oligonucleotides ranging from 10^2^ to 10^-4^ μM.

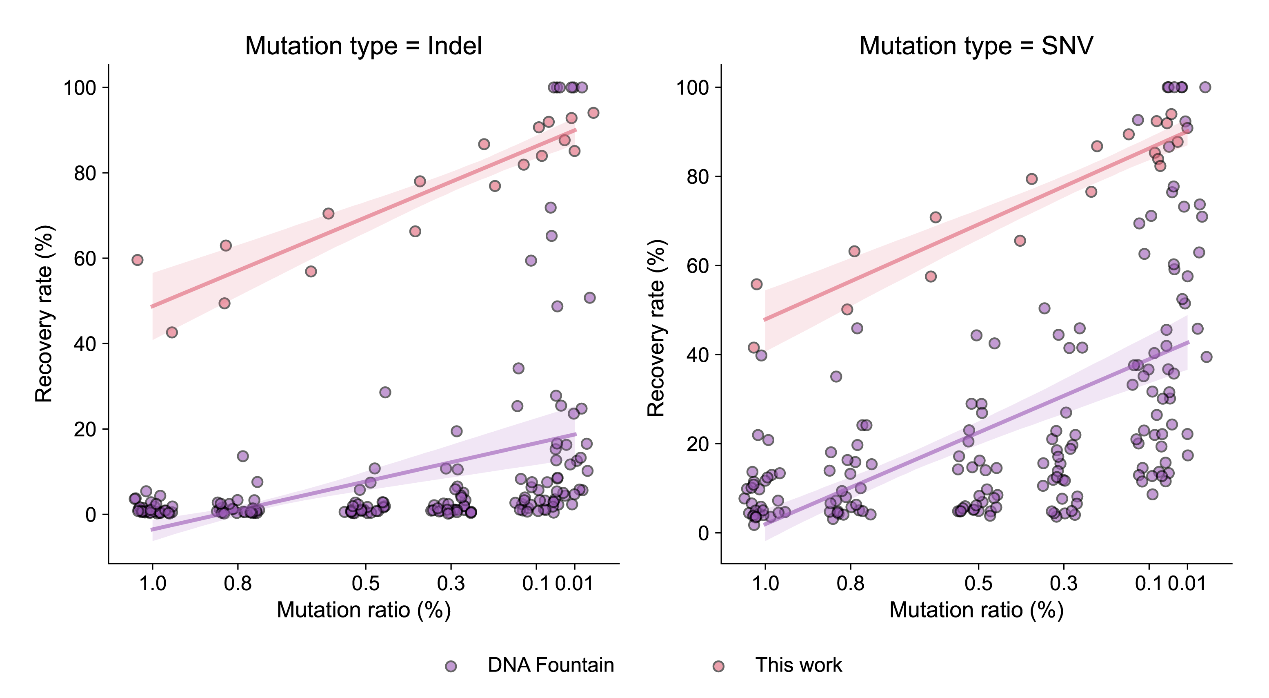

**Fig. S7.** Robustness characterization of DNA-SaM compared to Yin-Yang code and DNA fountain.

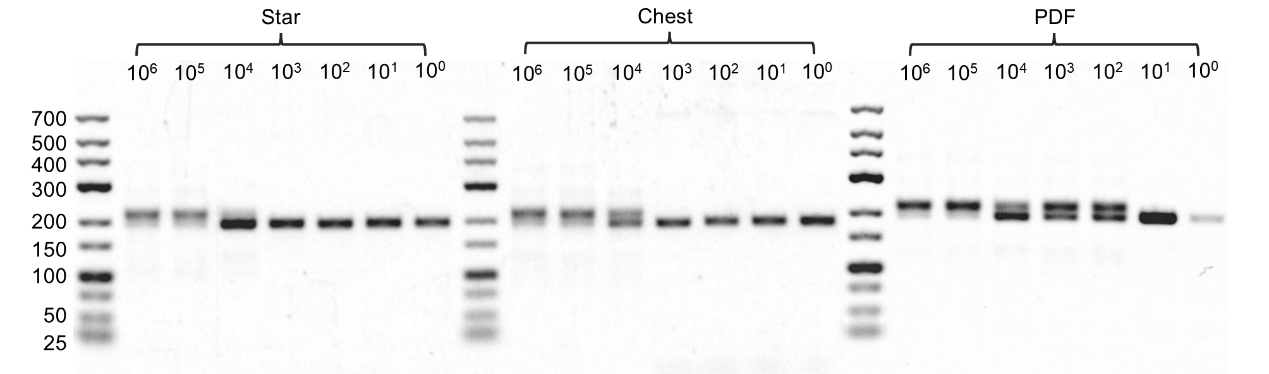

**Fig. S8.** Size verification after serial dilution PCR for oligo pools synthesized for three files. The oligo pools were diluted from 10^6^ to 10^0^ molecule copy numbers.

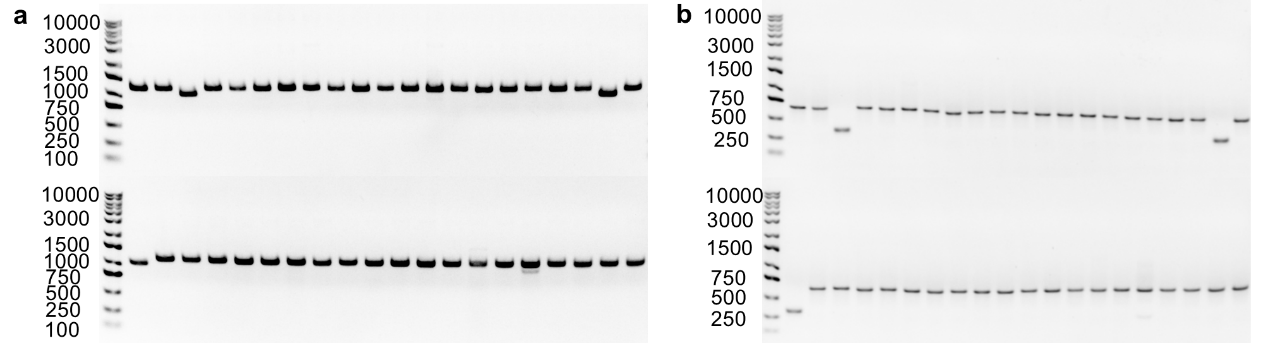

**Fig. S9.** Size verification performed by gel electrophoresis. **(a)** Verification of 41 strands of information-encoded DNA sequences inserted in plasmids. **(b)** Verification of 41 strands of information-encoded DNA sequences.

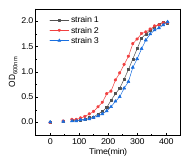

**Fig. S10.** Growth curves of three randomly selected *E.coli* strains containing information-encoded DNA sequences.

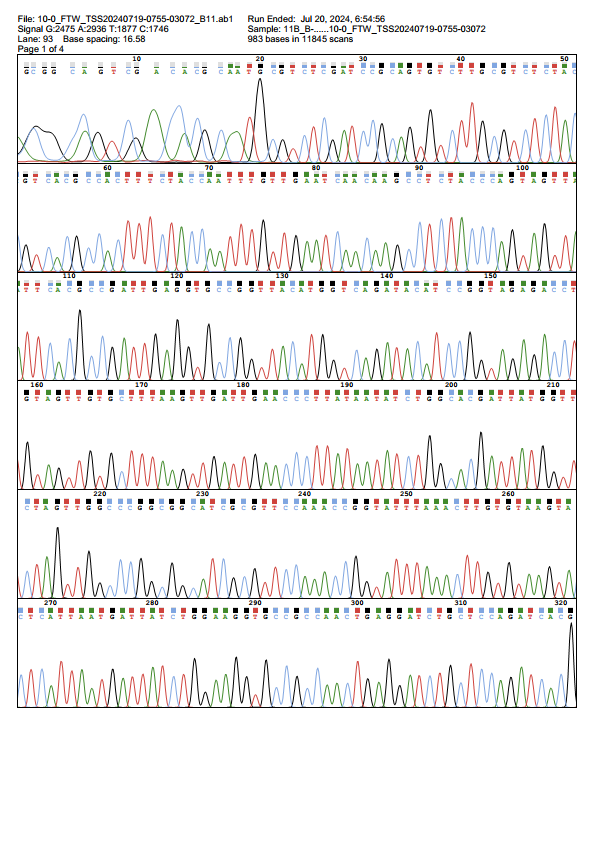

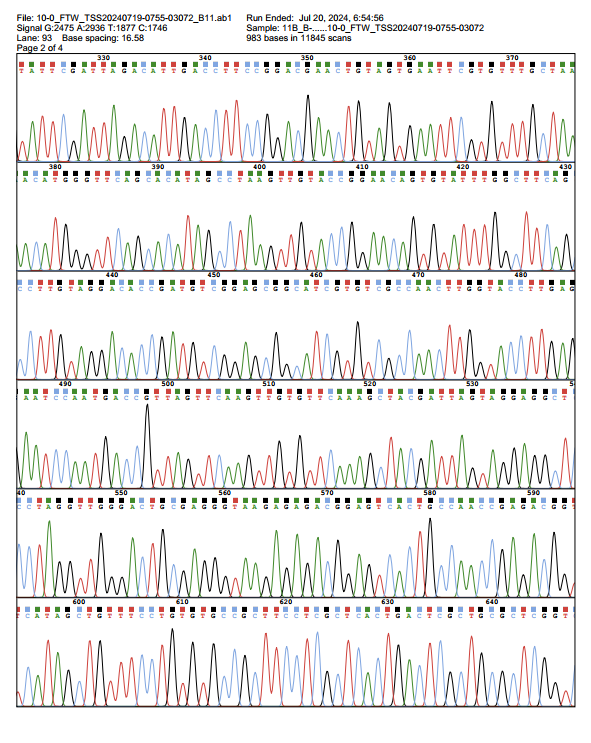

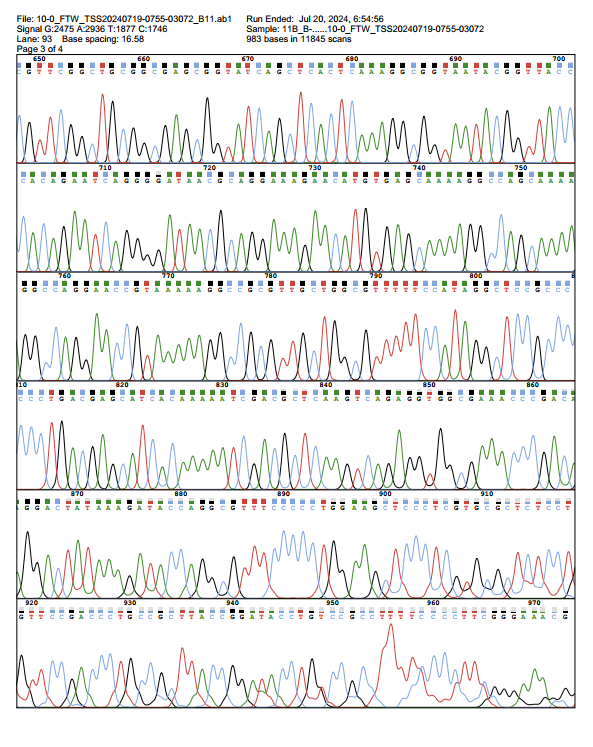

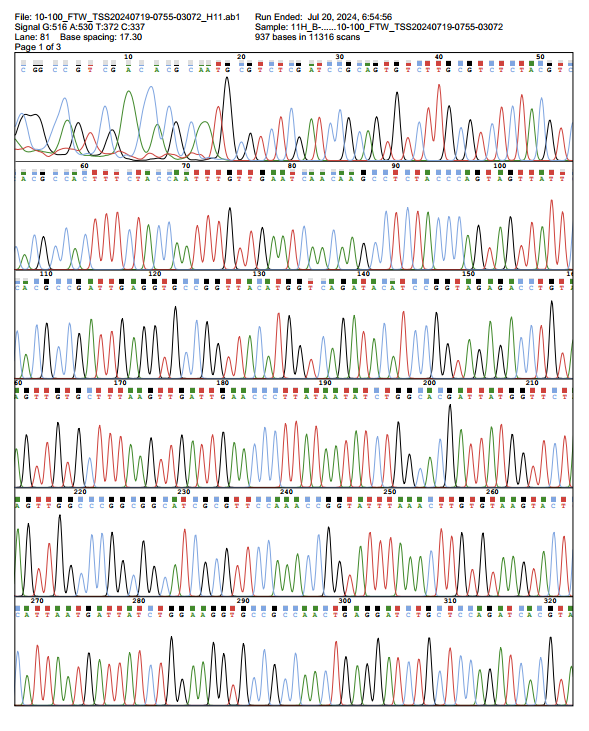

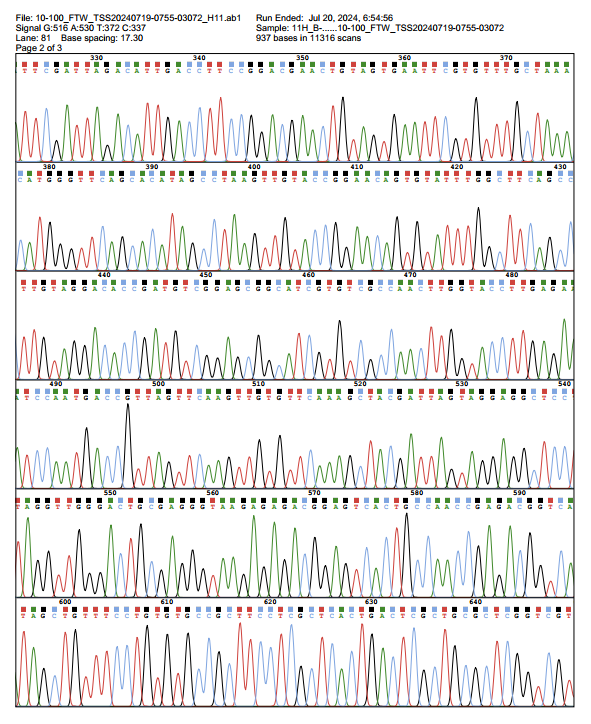

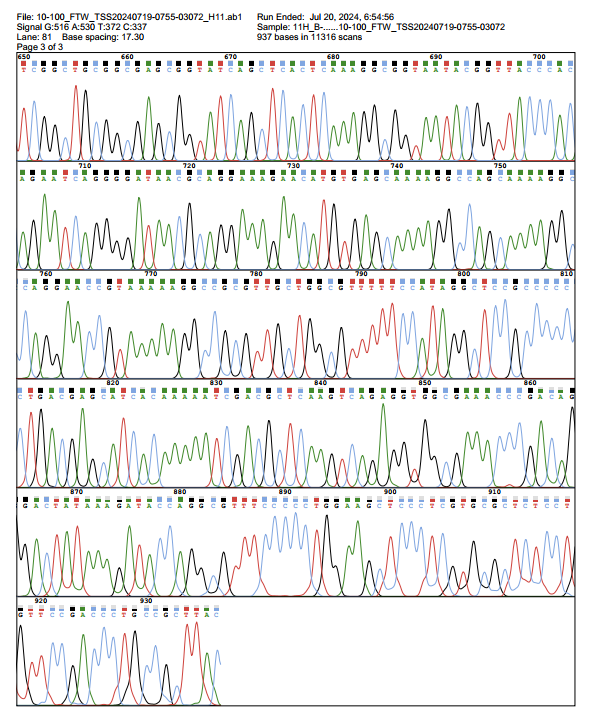

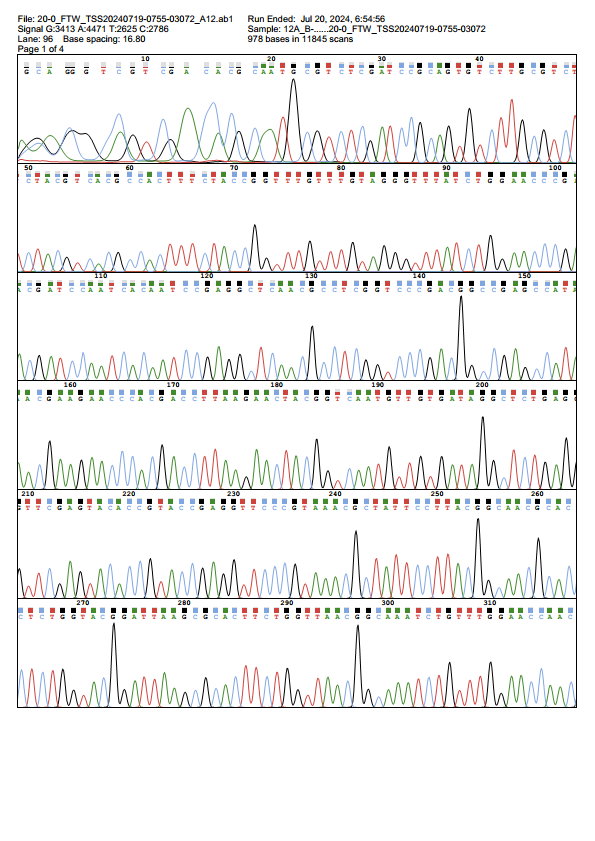

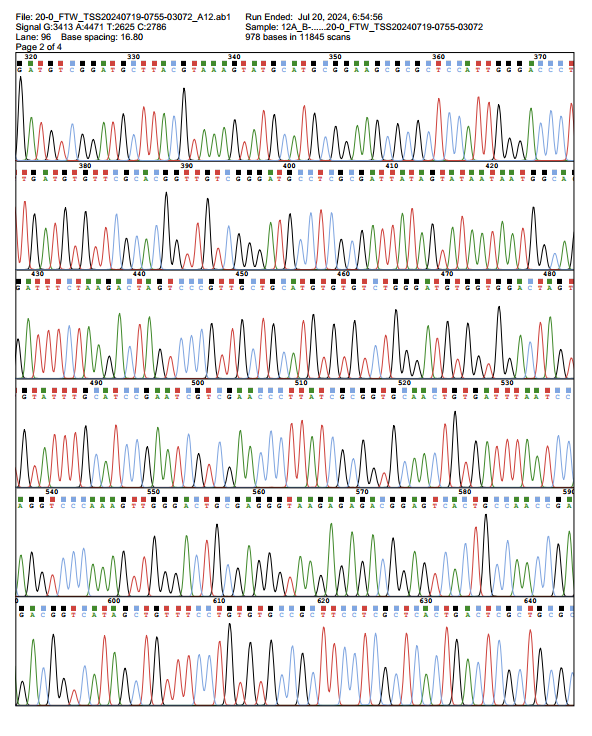

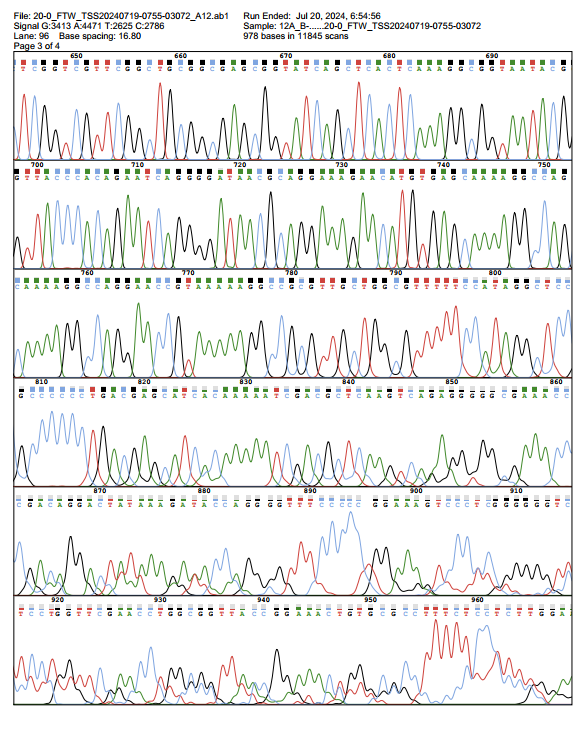

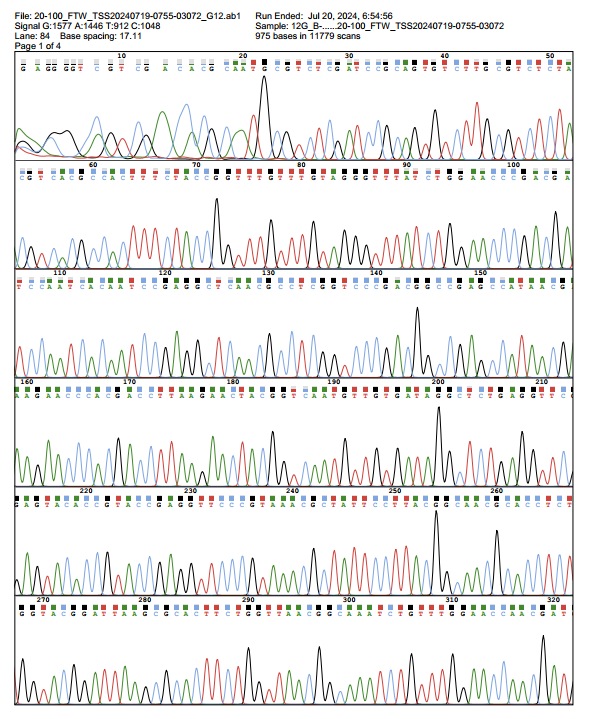

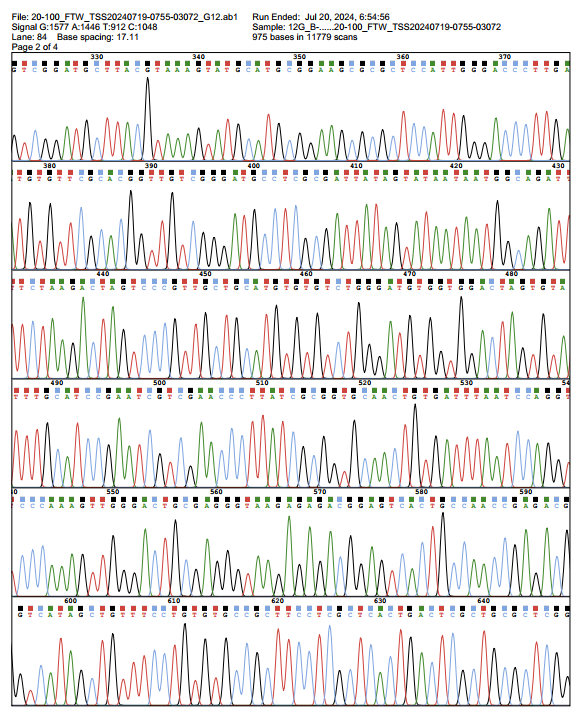

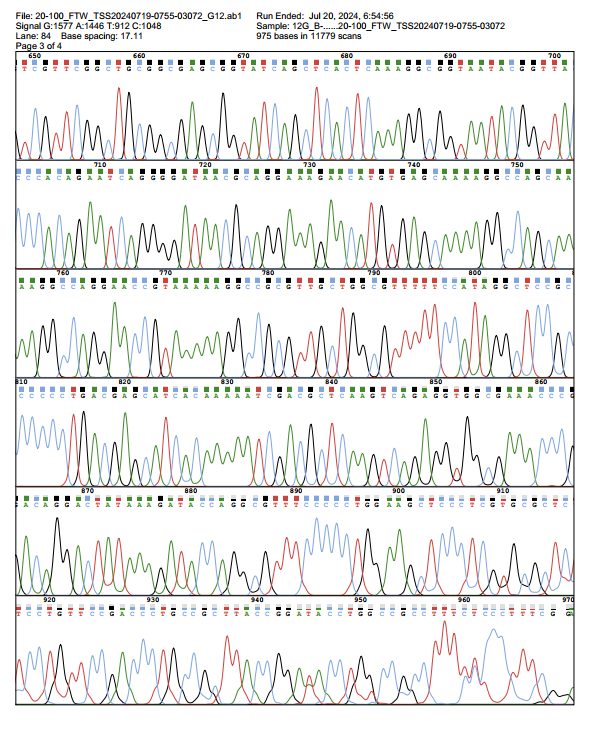

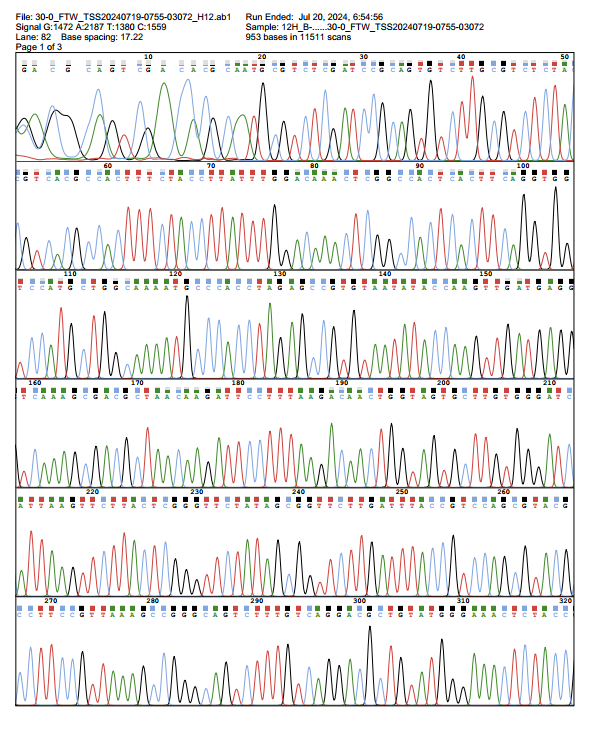

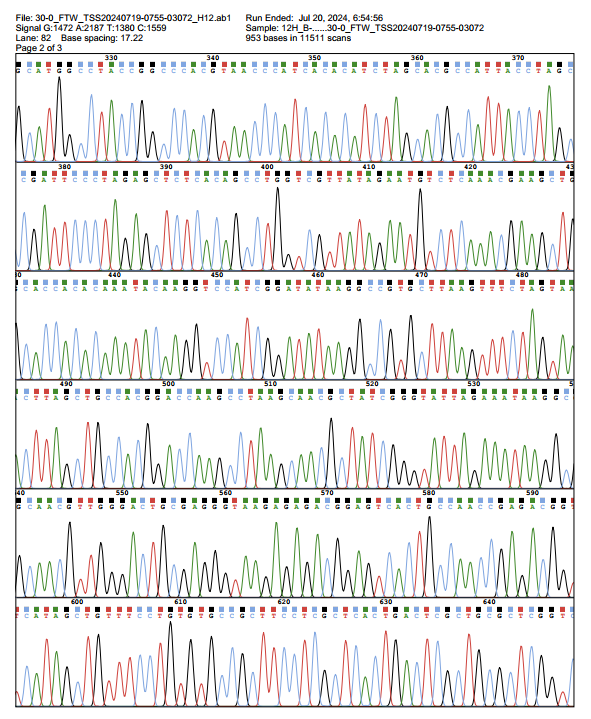

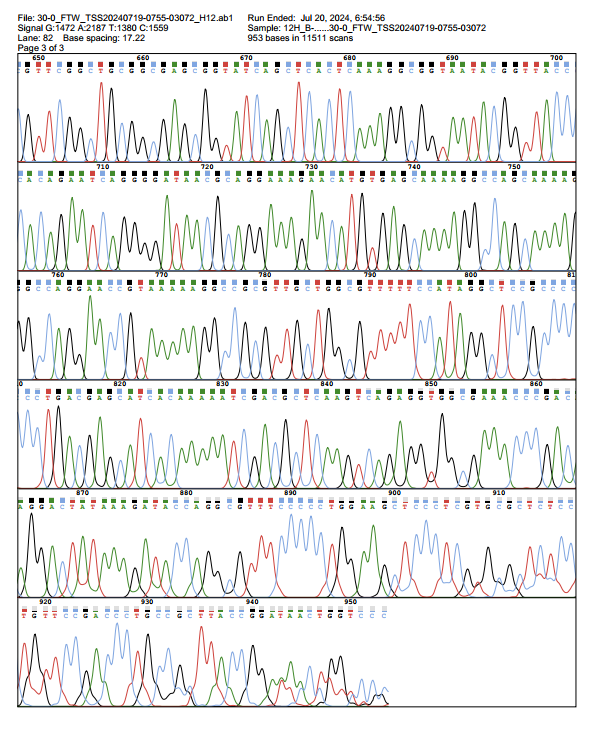

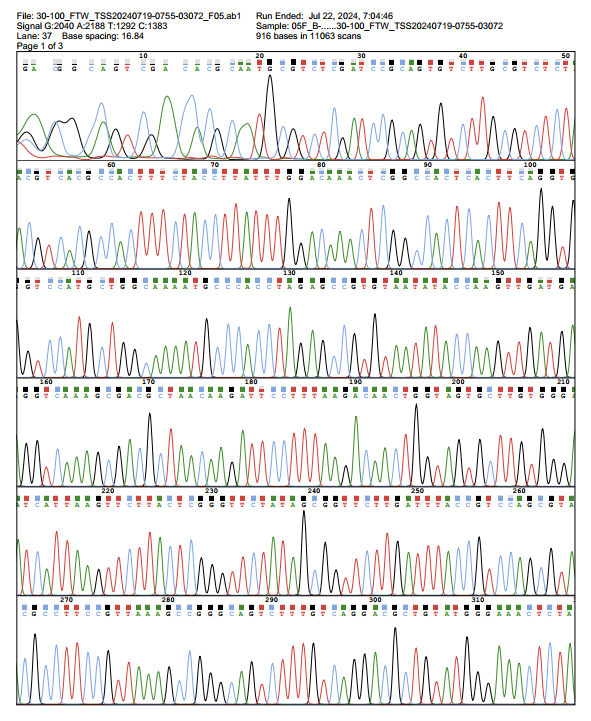

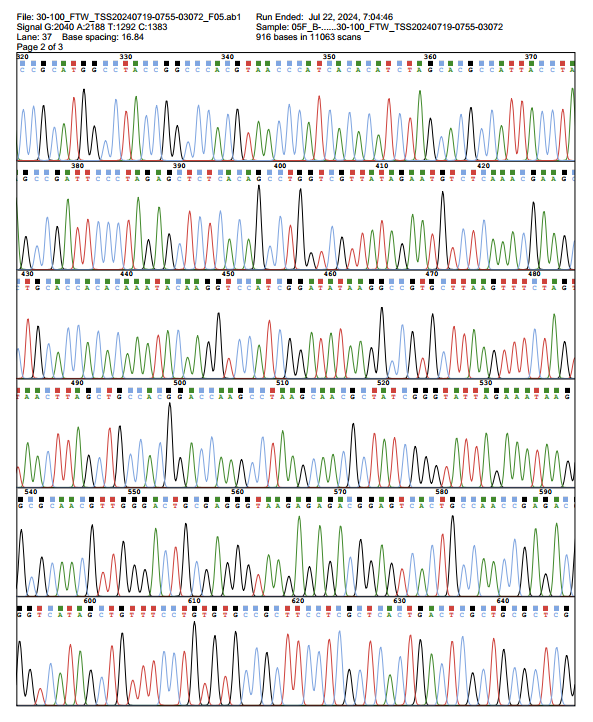

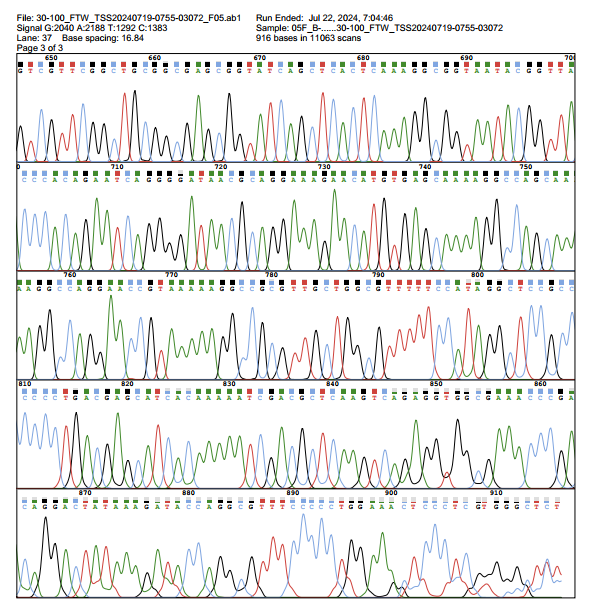

**Fig. S11.** Sequencing result for randomly selected three strains after culturing 0 and 100 generations.

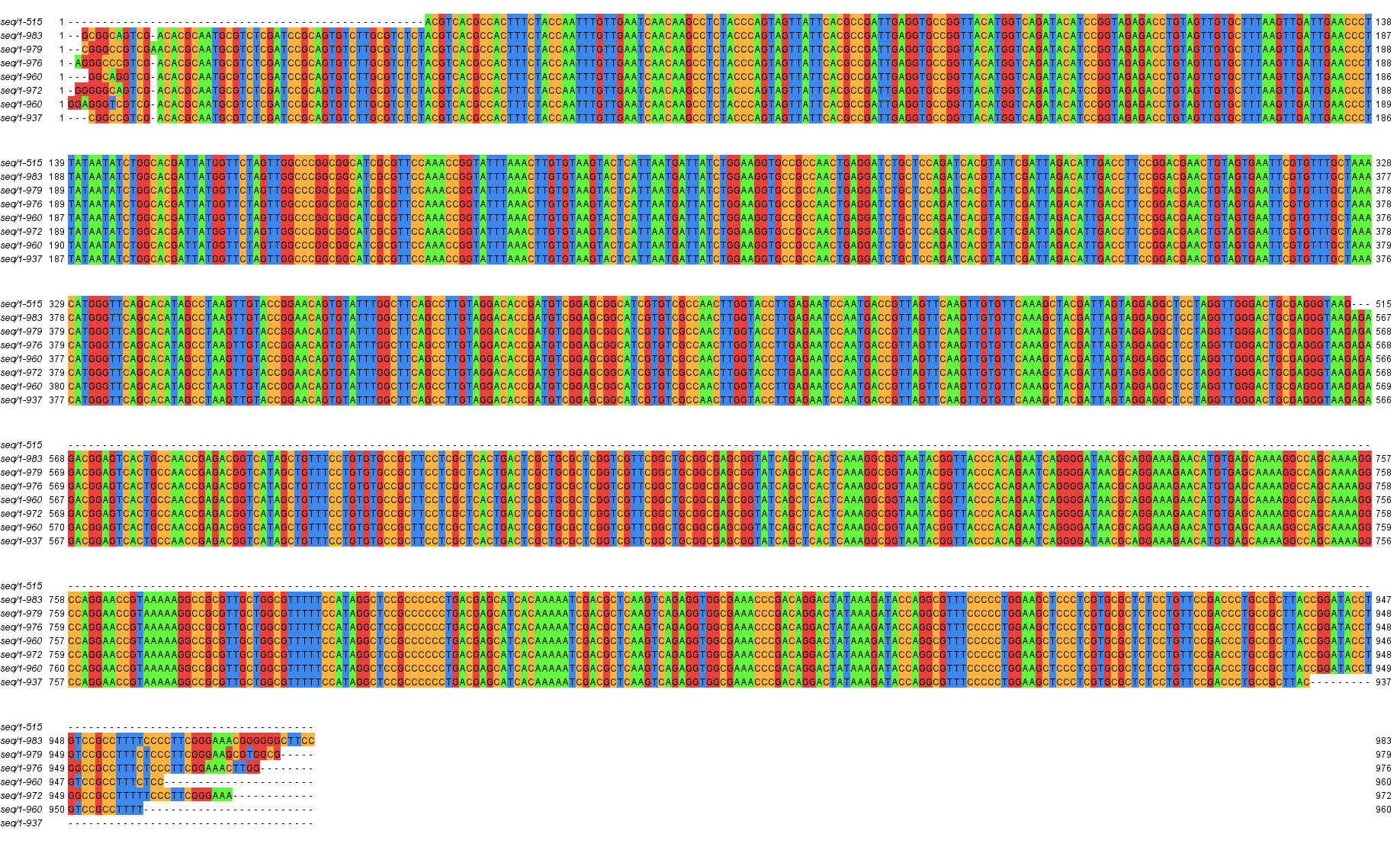

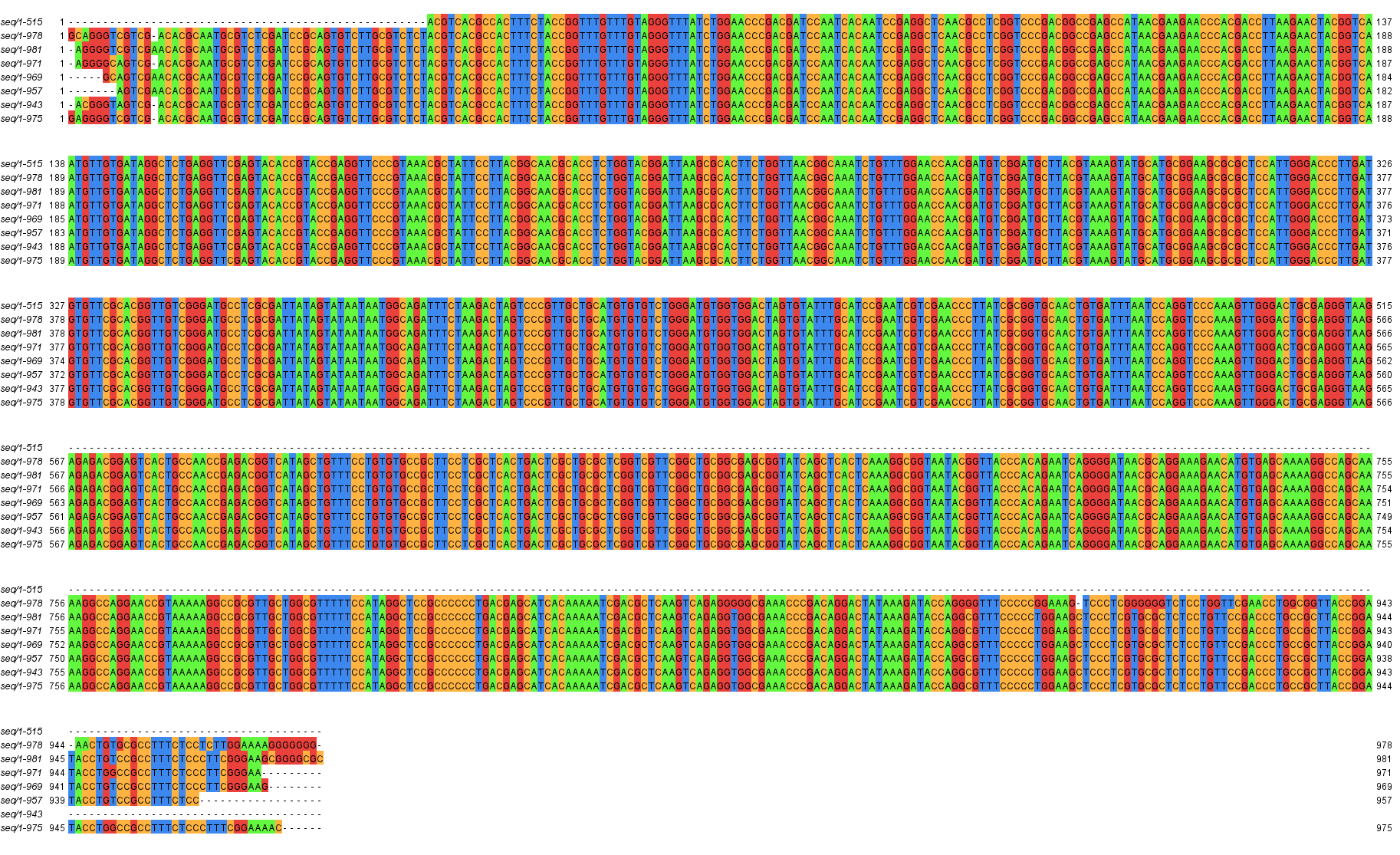

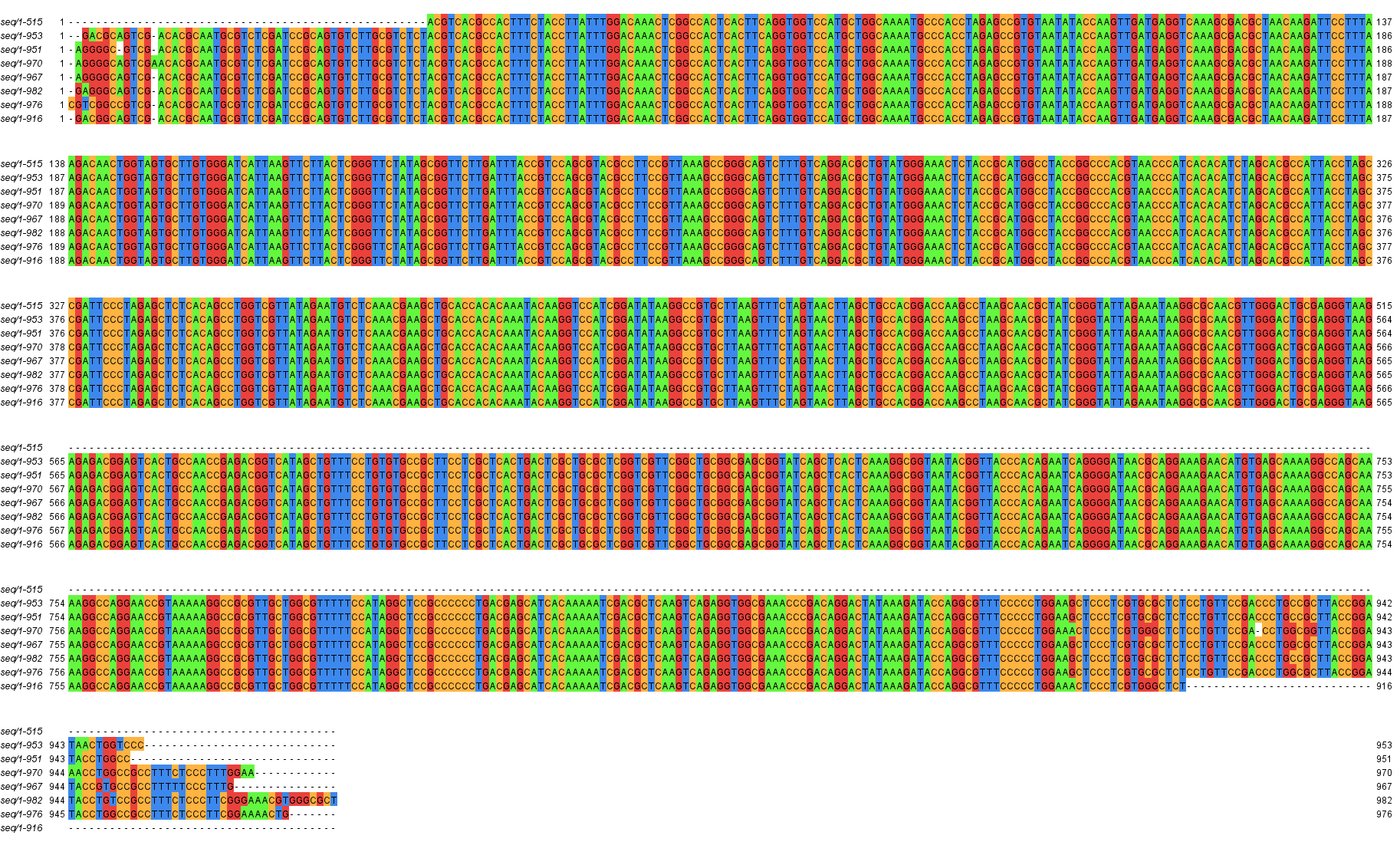

**Fig. S12.** Sequence alignment for the randomly selected three strains after culturing 0, 10, 20, 40, 60, 80 and 100 generation.

**Supplementary Tables**

**Table S1.** File type encoding table.

| **Icon** | **File type** | **File suffix** | **DNA** |
| --- | --- | --- | --- |
| 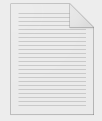 | Text file | .txt | ATCG |
|  |  | .doc/.docx | ATGC |
|  |  | .log | TACG |
| 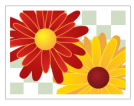 | Image file | .jpg/.jpeg | TCGA |
|  |  | .png | ACTG |
|  |  | .gif | ACGT |
|  |  | .bmp | AGCT |
|  |  | .tiff | AGTC |
| 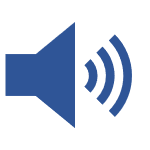 | Audio files | .mp3 | TAGC |
|  |  | .wav | TCAG |
|  |  | .aac | TGAC |
|  |  | .flac | TGCA |
|  |  | .ogg | CATG |
| 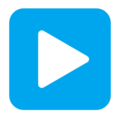 | video file | .mp4 | ATCA |
|  |  | .avi | ATAC |
|  |  | .mov | TCGC |
|  |  | .wmv | ACGC |
|  |  | .mvkv | AGGT |
| 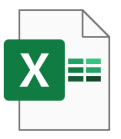 | Spreadsheet | .xls/.xlsx | CAGT |
|  |  | .csv | CTAG |
| 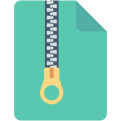 | Compress file | .zip | CTGA |
|  |  | .rar | CGAT |
|  |  | .7z | CGTA |
|  |  | . gz | GATC |
| 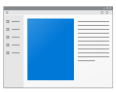 | Executable file | .exe | GACT |
|  |  | .app | GTCA |
| 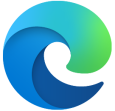 | Web page file | .html/.htm | GTAC |
|  |  | .css | GCTA |
| 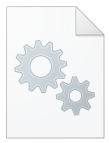 | Database file | .db | GATG |
|  |  | .sql | GTAG |
| 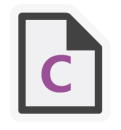 | Program source Source code file | .c/.cpp | CATC |
|  |  | .py | CTAC |
|  |  | .java | ACGA |
|  |  | .js | AGCA |
| 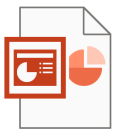 | Document files | .ppt | TCGT |
|  |  | .pdf | TGCT |
|  |  | .md | ACAC |
|  |  | .epub | CACA |

**Table S2.** DNA sequences used for PCR assembly after replacement.

| **Oligo no.^1^** | **Sequence (5’ to 3’)** | **Length (nt)** |
| --- | --- | --- |
| Replaced sequence | ACTCAGCCTCAGGTCGACACAATAAACCCTAGCAGTATGGGCTATTGGAGCTAGTACCGCCATTGTGGCGACGTGGGCGTGTCCGGCGCGGATAGGACACCTGGGAGCTCTAGGTTCTATGGGTGTTGCCGGCATGAATGGCTTACGAGGCCACCAGGGGATGCCAGATCCGATCGATCGATGCGACTTGCTGGACTCAGGTGAGTCGATCGAGTCGATGGAGCTGACAGAGCTGACTGTTAACCAGGTTAAGTAGGTCTAGTGTGTCTAGTGCGAATCCTAGAATCATTTCTGACAGCTGAATAATTGCCGCCAACAACCAGTCC | 326 |
| P’1 | ACTCAGCCTCAGGTCGACAC | 20 |
| P’2 | GGACTGGTTGTTGGCGGCAA | 20 |
| O’1 | ACTCAGCCTCAGGTCGACACAATAAACCCTAGCAGTATGGGCTATTG | 47 |
| O’2 | CCGGACACGCCCACGTCGCCACAATGGCGGTACTAGCTCCAATAGCCCATACTGCTAGG | 59 |
| O’3 | GTGGGCGTGTCCGGCGCGGATAGGACACCTGGGAGCTCTAGGTTCTATGGGTGTTGCCG | 59 |
| O’4 | CGGATCTGGCATCCCCTGGTGGCCTCGTAAGCCATTCATGCCGGCAACACCCATAGAAC | 59 |
| O’5 | GGGGATGCCAGATCCGATCGATCGATGCGACTTGCTGGACTCAGGTGAGTCGATCGAGT | 59 |
| O’6 | CTACTTAACCTGGTTAACAGTCAGCTCTGTCAGCTCCATCGACTCGATCGACTCACCTG | 59 |
| O’7 | GACTGTTAACCAGGTTAAGTAGGTCTAGTGTGTCTAGTGCGAATCCTAGAATCATTTCT | 59 |
| O’8 | GGACTGGTTGTTGGCGGCAATTATTCAGCTGTCAGAAATGATTCTAGGATTCGCA | 55 |

^1^: P’1 and P’2 stand for the primers for PCR assembly, O’1-8 are the oligonucleotides used for assembly.

**Table S3.** DNA sequences used for PCR assembly before replacement.

| **Oligo no.^2^** | **Sequence (5’ to 3’)** | **Length (nt)** |
| --- | --- | --- |
| Original sequence | ACTCAGCCTCAGGTCGACACAATAAACCCTAGAATCCTAGAATCCTAGAATCCTAGAATCCTAGAATCCTAGAATCCTAGAATCCTAGAATCCTAGAATCCTAGAATCCTAGAATCCTAGAATCCTAGAATCCTAGAATCCTAGAATCCTAGAATCCTAGAATCCTAGAATCCTAGAATCCTAGAATCCTAGAATCCTAGAATCCTAGAATCCTAGAATCCTAGAATCCTAGAATCCTAGAATCCTAGAATCCTAGAATCCTAGAATCCTAGAATCCTAGAATCCTAGAATCCTAGAATCCTAGAATCCTAGAATCCTAGAATCCTAGAATCCTAGAATCCTAGAATCCTAGAATCCTAGAATCCTAGAATCCTAGAATCCTAGAATCCTAGAATCCTAGAATCCTAGAATCCTAGAATCCTAGAATCCTAGAATCCTAGAATCCTAGAATCCTAGAATCCTAGAATCCTAGAATCATTTCTGACAGCTGAATAATTGCCGCCAACAACCAGTCC | 515 |
| P1 | ACTCAGCCTCAGGTCGACAC | 20 |
| P2 | TTGCCGCCAACAACCAGTCC | 20 |
| O1 | ACTCAGCCTCAGGTCGACACAATAAACCCTAGAATCCTAGAATCCTA | 47 |
| O2 | AGGATTCTAGGATTCTAGGATTCTAGGATTCTAGGATTCTAGGATTCTAGGATTCTAGG | 59 |
| O3 | TCCTAGAATCCTAGAATCCTAGAATCCTAGAATCCTAGAATCCTAGAATCCTAGAATCC | 59 |
| O4 | AGGATTCTAGGATTCTAGGATTCTAGGATTCTAGGATTCTAGGATTCTAGGATTCTAGG | 59 |
| O5 | TCCTAGAATCCTAGAATCCTAGAATCCTAGAATCCTAGAATCCTAGAATCCTAGAATCC | 59 |
| O6 | CTAGGATTCTAGGATTCTAGGATTCTAGGATTCTAGGATTCTAGGATTCTAGGATTCTA | 59 |
| O7 | AGAATCCTAGAATCCTAGAATCCTAGAATCCTAGAATCCTAGAATCCTAGAATCCTAGA | 59 |
| O8 | AGGATTCTAGGATTCTAGGATTCTAGGATTCTAGGATTCTAGGATTCTAGGATTCTAGG | 59 |
| O9 | TCCTAGAATCCTAGAATCCTAGAATCCTAGAATCCTAGAATCCTAGAATCCTAGAATCC | 59 |
| O10 | GATTCTAGGATTCTAGGATTCTAGGATTCTAGGATTCTAGGATTCTAGGATTCTAGGAT | 59 |
| O11 | AATCCTAGAATCCTAGAATCCTAGAATCCTAGAATCCTAGAATCCTAGAATCCTAGAAT | 59 |
| O12 | AGCTGTCAGAAATGATTCTAGGATTCTAGGATTCTAGGATTCTAGGATTCTAGGATTC | 58 |
| O13 | AGAATCATTTCTGACAGCTGAATAATTGCCGCCAACAACCAGTCC | 45 |

^2^: P1 and P2 stand for the primers for PCR assembly, O1-13 are the oligonucleotides used for assembly.

**Table S4.** Editing of bio-functional sequences in genome by SAM.

| **Gene_Name** | **Description** | **Sequence** |
| --- | --- | --- |
| TFAP2E | transcription factor AP-2 epsilon | CGCCTCAGGCG |
| CBFB | core-binding factor subunit beta | TCTGTGGTTTG |
| NRF1 | nuclear respiratory factor 1 | GCGCCTGCGCA |
| EMC8_1 | ER membrane protein complex subunit 8 | GTCGTAAAATGCCGGGCCGGGCTGCTTTCCGGCAGGCGTTGGAGAGGCTGCTTTCCGGCG |
| COX4I1_1 | cytochrome c oxidase subunit 4I1 | CGGGCGGAGTCTTCCTCGATCCCGTGGTGCTCCGCGGCGCGGCCTTGCTCTCTTCCGGTC |
| UBC_1 | ubiquitin C | CGTCACCCGTTCTGTTGGCTTATAATGCAGGGTGGGGCCACCTGCCGGTAGGTGTGCGGT |
| ­D20S16 | Human centromeric satellite | CAGCTCCACGACCCTCAAACTAGAACAAGACCTCTCCTCCCCGGGTCNCCAGCTNCCNGACCCTCGAACTACAACAACATTGCTCCTCCCTGGGTCTT |
| HSAT5 | Human centromeric satellite 5 | CACTGACCAGGNCCCCACTGACNAGGCCCCACTGACNAGGCCCCACTGACCAGGTCCCCACTGACNAGGCCTCACTGACCAGGCCT |
| MADE1 | MAriner Derived Element 1, a TcMar-Mariner DNA transposon | TTAGGTTGGTGCAAAAGTAATTGCGGTTTTTGCCATTACTTTTAATGGCAAAAACCGCAATTACTTTTGCACCAACCTAA |
| D22Z3 | A novel centromeric repetitive DNA from human chromosome 22 | TTCCAGAACACTGCTGCTGGGNTCTGAATGTTTGTCCCTCACATAGGATTCCAGAACACTGCTGCTGGGNTCTGAGTGTTTGTCCCTCACATAGGA |
| LOC | A gene group involved in lactose breakdown | ……TGTTGT……ACAACA…… |
| Rho | Transcription termination protein | ...GCGAAG...CUUCGC... |
| Ara | breakdown metabolism of synthetic sugars | ……TTGACA……AACTGT…… |

**Table S5.** Comparison of performance guaranteed by various algorithms when encoding text, image, audio, and video files.

| **File type** | **Algorithm** | **Encoding sequence number** | **TATA box** | **SD sequence** | **GAAT box** | **Tandem repeat** |
| --- | --- | --- | --- | --- | --- | --- |
| Text | DNA-Aeon | 189 | 37 | 156 | 492 | 0 |
|  | Goldman et al. | 432 | 0 | 0 | 0 | 77 |
|  | YYC | 86 | 0 | 0 | 0 | 3 |
|  | This work | 357 | 0 | 0 | 0 | 0 |
| Figure | DNA-Aeon | 987 | 28 | 124 | 418 | 0 |
|  | Goldman et al. | 2313 | 0 | 0 | 0 | 269 |
|  | YYC | 462 | 0 | 0 | 0 | 26 |
|  | This work | 1878 | 0 | 0 | 0 | 0 |
| Audio | DNA-Aeon | 469 | 51 | 225 | 700 | 0 |
|  | Goldman et al. | 1096 | 0 | 0 | 0 | 279 |
|  | YYC | 287 | 0 | 0 | 0 | 153 |
|  | This work | 2220 | 0 | 0 | 0 | 0 |
| Video | DNA-Aeon | 1167 | 87 | 336 | 1180 | 1 |
|  | Goldman et al. | 2736 | 0 | 0 | 0 | 2127 |
|  | YYC | 644 | 0 | 0 | 0 | 224 |
|  | This work | 2220 | 0 | 0 | 0 | 0 |

A 10,683-byte text, 56,275-byte image, 26,660-byte audio, 66,518-byte video for encoding by DNA-Aeon, Goldman *et al.*, and YYC. The computer environment: CUP: Inter® Core ™ i7-10700 CPU @2.90 GHZ, RAM, 16 GB, at a Microsoft X64 system.

**Table S6.** Operation time and times of forbidden sequence replacement for various numbers and types of sequences.

| **Length of forbidden sequence** | **Time (s)** | **Number of forbidden sequences** | **Replace times** |
| --- | --- | --- | --- |
| 8 | 5.812778 | 20 | 582 |
| 12 | 5.377404 | 20 | 24 |
| 16 | 5.271085 | 20 | 20 |
| 20 | 5.259195 | 20 | 20 |
| 24 | 5.269698 | 20 | 20 |
| 28 | 5.276229 | 20 | 20 |
| 32 | 5.288354 | 20 | 20 |
| 36 | 5.266013 | 20 | 20 |
| 40 | 5.238246 | 20 | 20 |
| 44 | 5.254269 | 20 | 20 |

**Table S7.** Time consumption for encoding various sizes of data by algorithms including Goldman *et al.*^1^, YYC ^2^, DNA fountain ^3^, DNA-Aeon ^4^, and this work. Unit of time: second.

| **File size (byte)** | **Goldman et al.** | **YYC** | **DNA fountain** | **DNA-Aeon** | **This work** |
| --- | --- | --- | --- | --- | --- |
| 10683 | 0.483433962 | 0.254195309 | 3.159500837 | 2.566871405 | 0.022252083 |
| 275544 | 11.95363593 | 7.118596387 | 978.3797209 | 5.707934141 | 0.629515171 |
| 625899 | 27.67560244 | 27.61254721 | 2289.543312 | 7.368292332 | 1.40004158 |
| 886155 | 39.31600904 | 52.39429502 | 3498.153245 | 14.27031469 | 1.962709188 |
| 1788217 | 103.3991609 | 435.6180344 | 10063.88924 | 24.59978914 | 3.448956013 |
| 5518673 | 324.4636273 | 3835.422735 | 66886.64121 | 126.0928905 | 10.42022204 |
| 6246512 | 385.1618769 | 4627.24138 | 83879.89958 | 186.0592549 | 11.89367843 |
| 7853393 | 465.0948634 | 6442.883514 | 128346.9829 | 254.7600585 | 15.0333178 |
| 8608227 | 508.2281826 | 7657.267165 | 155823.8981 | 335.1617985 | 16.38237405 |

**Table S8.** Time consumption for encoding files in GB, TB, and PB level by algorithms including Goldman *et al.*^1^, YYC ^2^, DNA fountain ^3^, DNA-Aeon ^4^, and this work. Unit of time: second.

| **File size** | **DNA fountain** | **YYC** | **DNA-Aeon** | **Goldman et al.** | **This work** |
| --- | --- | --- | --- | --- | --- |
| GB | 6.51E+01 | 2.48E+00 | 1.59E-01 | 2.05E-03 | 6.43E-05 |
| TB | 6.81E+07 | 2.59E+06 | 1.67E+05 | 2.10E+00 | 6.58E-02 |
| PB | 7.14E+13 | 2.72E+12 | 1.75E+11 | 2.15E+03 | 6.74E+01 |

**Table S9.** File encoding details for experimental verification.

| **File name** | **File size** | **Oligo number** | **File permission** | **File primer (left)** | **File primer (right)** | **File random address** |
| --- | --- | --- | --- | --- | --- | --- |
| star.mp4 | 243,517 | 9744 | user | ATTGCCCCTTGGACCCAACG | CTCATCTCCGGCAGCAGTAG | [1009, 488, 12, 720, 929, 365, 771, 586, 871, 577, 605] |
| chest_x_ray.png | 266,208 | 10651 | admin | TCGCCCGAAGAAAAGACTCC | TACGAACGCAGGGTGTAAGG | [387, 499, 117, 404, 374, 920, 141, 732, 214, 560, 330] |
| The_Google_File_System.pdf | 275,544 | 11024 | admin | GCCACAGATTTCGCGATACC | GGAGTCATGTGGGGCCTTTC | [399, 372, 308, 649, 25, 744, 365, 350, 672, 937, 490] |

**Table S10.** Oligo pools designed for proof-of-concept experiments (Provided in Supplementary material 2).

**Table S11.** DNA sequences designed for storing A very short story by Ernest Miller Hemingway (Provided in Supplementary material 3).

**Table S12.** Primers for PCR amplification of oligo pools generated from Star, Chest, and PDF.

| **Oligo no.** | **Sequence (5’ to 3’)** | **Length (nt)** |
| --- | --- | --- |
| P1-star | ATTGCCCCTTGGACCCAACG | 20 |
| P2-star | CTACTGCTGCCGGAGATGAG | 20 |
| P1-chest | TCGCCCGAAGAAAAGACTCC | 20 |
| P2-chest | CCTTACACCCTGCGTTCGTA | 20 |
| P1-PDF | GCCACAGATTTCGCGATACC | 20 |
| P2-PDF | GAAAGGCCCCACATGACTCC | 20 |

1	Goldman, N. *et al.* Towards practical, high-capacity, low-maintenance information storage in synthesized DNA. *Nature* **494**, 77-80, (2013).

2	Ping, Z. *et al.* Towards practical and robust DNA-based data archiving using the yin-yang codec system. *Nat. Comput. Sci.* **2**, 234-242, (2022).

3	Erlich, Y. & Zielinski, D. DNA fountain enables a robust and efficient storage architecture. *Science* **355**, 950-954, (2017).

4	Welzel, M. *et al.* DNA-aeon provides flexible arithmetic coding for constraint adherence and error correction in DNA storage. *Nature Communications* **14**, (2023).
